# Emergence and maintenance of variable-length actin filaments in a limiting pool of building blocks

**DOI:** 10.1101/2021.11.07.467615

**Authors:** Deb Sankar Banerjee, Shiladitya Banerjee

## Abstract

Actin is one of the key structural components of the eukaryotic cytoskeleton that regulates cellular architecture and mechanical properties. Dynamic regulation of actin filament length and organization is essential for the control of many physiological processes including cell adhesion, motility and division. While previous studies have mostly focused on the mechanisms controlling the mean length of individual actin filaments, it remains poorly understood how distinct actin filament populations in cells maintain different lengths using the same set of molecular building blocks. Here we develop a theoretical model for the length regulation of multiple actin filaments by nucleation and growth rate modulation by actin binding proteins in a limiting pool of monomers. We first show that spontaneous nucleation of actin filaments naturally leads to heterogeneities in filament length distribution. We then investigate the effects of filament growth inhibition by capping proteins and growth promotion by formin proteins on filament length distribution. We find that filament length heterogeneity can be increased by growth inhibition, whereas growth promoters do not significantly affect length heterogeneity. Interestingly, a competition between filament growth inhibitors and growth promoters can give rise to bimodal filament length distribution as well as a highly heterogeneous length distribution with large statistical dispersion. We quantitatively predict how heterogeneity in actin filament length can be modulated by tuning F-actin nucleation and growth rates in order to create distinct filament subpopulations with different lengths.

**SIGNIFICANCE:** Actin filaments organize into different functional network architectures within eukaryotic cells. To maintain distinct actin network architectures, it is essential to regulate the lengths of actin filaments. While the mechanisms controlling the lengths of individual actin filaments have been extensively studied, the regulation of length heterogeneity in actin filament populations is not well understood. Here we show that the modulation of actin filament growth and nucleation rates by actin binding proteins can regulate actin length distribution and create distinct sub-populations with different lengths. In particular, by tuning concentrations of formin, profilin and capping proteins, various aspects of actin filament length distribution can be controlled. Insights gained from our results may have significant implications for the regulation of actin filament length heterogeneity and architecture within a cell.

## INTRODUCTION

In eukaryotic cells, actin filament growth and turnover are tightly regulated for coordinating a diverse set of physiological processes including cell motility (1–3), protrusion formation (4–6), endocytosis (7), wound healing (8, 9), synaptic activity (10), cytokinesis (11, 12), and embryonic development (13–16). These different functions often require distinct subpopulations of actin filaments organized into different lengths and architectures (17). Study of length control mechanisms of intracellular structures has been an active area of research. Traditionally, these studies mostly focus on understanding the physical principle and molecular mechanism that give rise to structures of a typical length (18–23), with a relatively narrow length distribution. The origin of length heterogeneity and the mechanisms to create and maintain a heterogeneous population is not well understood. Here we study mechanisms of controlling heterogeneity in length of a population of actin filaments.

The molecular processes underlying actin filament growth have been extensively studied both theoretically and experimentally. Existing models for actin filament growth can be categorized into two main classes according to the availability of monomers (24): growth in an open system where the monomer concentration remains unchanged in time, and growth in a closed system where free monomer concentration decreases as filaments increase in length (keeping the total amount of monomers conserved). Length control mechanisms have been proposed for actin filaments in both the above cases to understand how filaments of a typical length can be achieved (22, 25–27). But, how filaments of different lengths can be obtained is much less understood (28). This is relevant for a cell where actin filaments are found in a wide diversity of lengths. In this study we consider actin filament growth in a closed system, where the monomer pool is limited (29). Though it is not well established if *in vivo* actin systems can be considered to be assembled from a limiting monomer pool, recent studies show that different actin structures in a cell often compete for the same limiting pool of monomeric actin (30, 31). In this study, we use theory and simulations to demonstrate how actin filament length heterogeneity can be regulated and we demonstrate how distinct actin filament subpopulations may emerge in a limiting pool of monomers and actin binding proteins, specifically formin, profilin and capping proteins. Since we limit our study to interactions between actin and only a few actin regulatory proteins, the results from our theoretical model can be best captured with an *in vitro* experimental system, where the concentrations of actin and its regulatory proteins can be precisely controlled.

Previous theoretical studies have shown that nucleation and polymerization of actin filaments in a limiting monomer pool results in an exponential length distribution at steadystate (24, 32–34), with the standard deviations in length fluctuations as large as the mean. The timescale for reaching such a steady-state length distribution is found to be *>* 3 hours *in vitro* (32, 35), while simulations predicted the timescale to be on the order of days (26, 33, 34) under physiological conditions (e.g., cell volume ∼ 1000 *µm*^3^ and total monomer concentration 20 − 100 *µ*M), making the steady-state length distribution less relevant for many of the physiological processes that occur within a timescale of minutes to hours. It has been previously shown that the dynamics of actin length distribution can be segregated into a fast regime of nucleation and growth, and a slow regime of length rearrangement between actin filaments via exchanging monomers (33). In the initial regime, the length distribution dynamics has a convective nature (mean of the distribution increases), while in the later regime the filament lengths undergo a diffusive dynamics (variance increases keeping mean constant) (33) (Fig S1). We are, however, interested in actin length distribution during an intermediate regime where the number of filaments and the mean filament length has reached a steady state. This intermediate regime is attained within minutes and the resulting length distribution remains approximately invariant over a timescales of hours, making it physiologically relevant as many actin structures such as filopodia, lamellipodia turns over within or before an hour (36–38). In this regime, the growth of the actin filament population can be regulated by both spontaneous filament nucleation and actin binding proteins such as formin, profilin and capping proteins. It is important to note that actin filament turnover by spontaneous fragmentation, annealing, and cofilin-mediated severing may affect the long-time actin dynamics in an actin concentrationdependent manner, thereby altering the time evolution of filament length distribution. Here we do not consider such effects and restrict our study to smaller actin concentrations, focusing on timescales short compared to the timescales over which fragmentation or annealing effects become prominent (see Methods).

We show that spontaneous nucleation of actin filaments and interactions with specific actin binding proteins (ABPs) can induce significant heterogeneity in actin filament length, persisting for several hours. We specifically study the effects of growth promoting ABPs like formin and growth inhibiting ABPs like capping proteins on the regulation of actin filament length distribution. We show that formin and capping protein concentrations can be tuned to regulate the heterogeneity in actin filament length in the filament population. In particular, we find that formin and capping proteins, when present in similar concentrations, can give rise to a bimodal length distribution that represent two distinct subpopulations of long and short actin filaments.

## METHODS

### Computational model

We first describe the computational model for the nucleation and growth of actin filaments in the presence of actin binding proteins. All the chemical reactions involved in F-actin growth and nucleation are listed in Fig. 1, with the parameter values given in Table. 1. We use a stochastic reaction network framework based on Gillespie’s first algorithm (39) to simulate the length dynamics of actin filaments. Spontaneous nucleation of actin filaments involves sequential formation of polymerization intermediates, actin dimers and trimers, with the trimers acting as the seed for nucleation (40–42) (Reactions 2-5 in Fig. 1 and Fig. 2a). We consider the growth of filamentous actin (F-actin) via association and dissociation of globular actin (G-actin) from the barbed end (Reaction 6 in Fig. 1 and Fig. 2a), with the association and dissociation rates *K*^on^ and *K*^off^ (see Supplemental Methods). For simplicity, we do not consider the nucleotide state of the monomers in the filament and assume that free monomers in the pool are ATP-bound.

**Figure 1:**
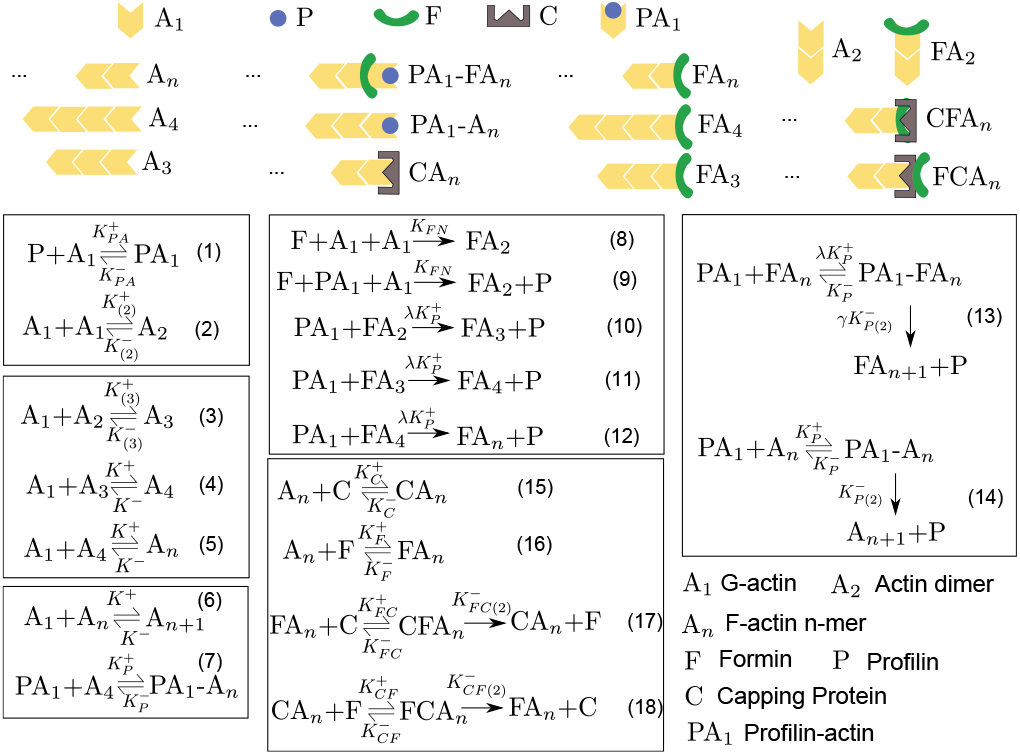
Model schematic. Schematic diagram showing all the chemical species and reactions involving the F-actin nucleation and growth in a limiting G-actin pool. Here *A*_1_, *A*_2_, …, *A*_*n*_ denote actin monomer, dimer and *n*-mer filaments (*n >* 4), respectively. The assembly and disassembly rates are defined in the schematic with the accompanying reaction numbers. The abbreviation XA_*n*_ stands for a filament with X-bound barbed end and PA_1_-X stands for profilin-actin monomer bound to the X species. FCA_*n*_ and CFA_*n*_ denote the two possible cases of the “decision-complex” formed via subsequent capping and formin (or formin and capping for CFA_*n*_) binding to a barbed end. The corresponding rate constants and parameters are listed in Tables 1 and Table 2.

**Figure 2:**
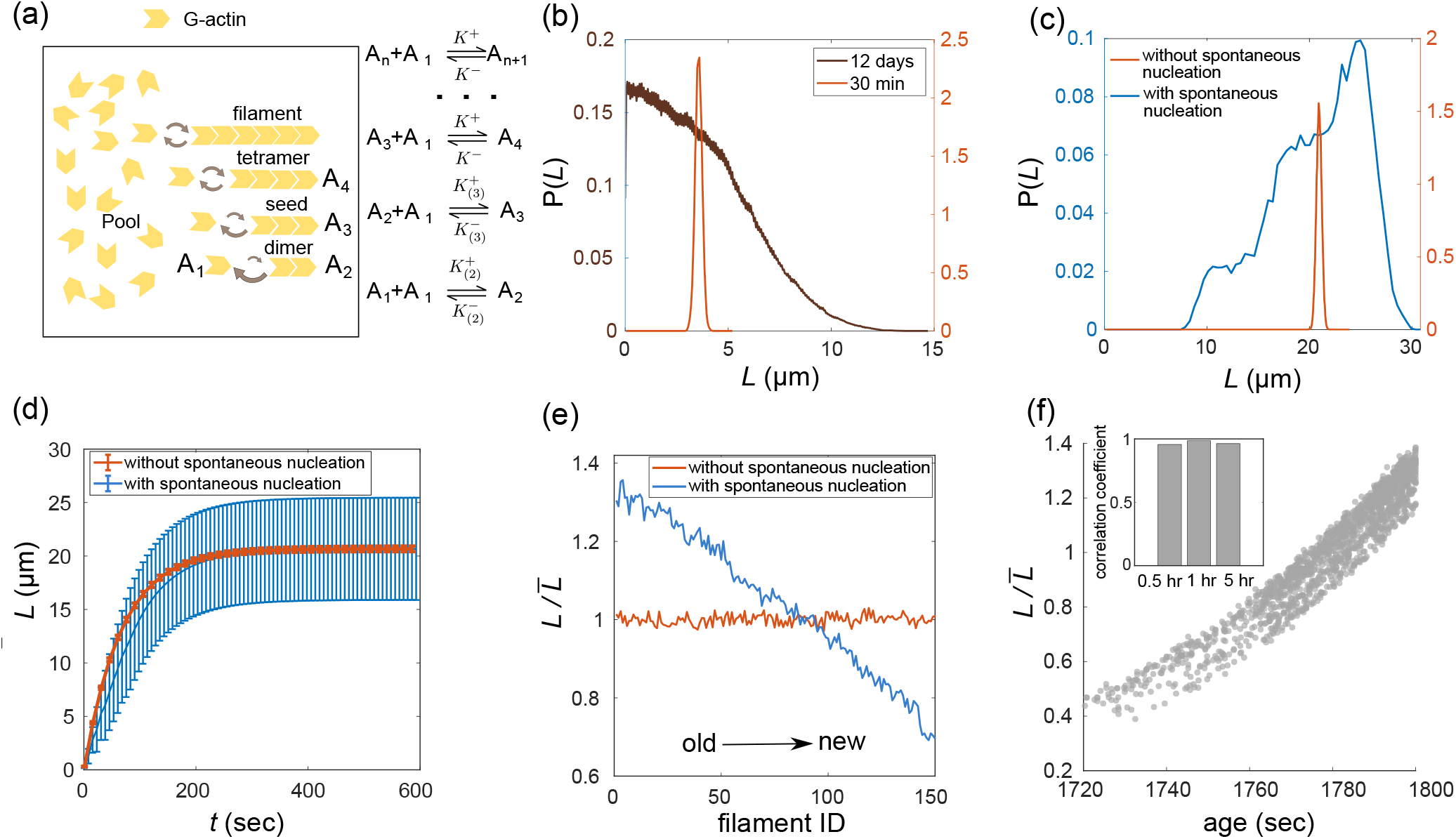
Spontaneous nucleation promotes heterogeneity in actin filament lengths. (a) Schematic diagram of spontaneous nucleation and growth of F-actin in a limiting G-actin pool. Here *A*_1_, *A*_2_, …, *A*_*n*_ stand for actin monomer, dimer and *n*-mer filaments, respectively. The assembly and disassembly rates are defined by the accompanying reaction diagrams. (b) Length distribution of actin filaments growing in a limiting monomer pool without spontaneous nucleation, showing a narrow length distribution at *t* = 30 min (red), and exponential-like distribution at *t* = 12 days. (c) The length dynamics in presence of spontaneous nucleation (blue) leads to a significantly broader length distribution as compared to growth of fixed number of F-actin from a limiting monomer pool (red). The broader length distribution originates from the underlying heterogeneity in filament length. (d) Time evolution of the mean filament length, with (blue) and without (red) spontaneous nucleation. (e) Length of filaments, ordered according to their age with ’0’ being the oldest, i.e. the first filament nucleated. The result is presented in terms of length (*L*) divided by mean filament length (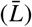). There is negligible length heterogeneity in filament population growing without spontaneous nucleation (red) as compared to filaments growing with spontaneous nucleation (blue). (f) Filament length and age are highly correlated in presence of spontaneous nucleation. (inset) The emergent correlation is long-lived and remains almost unaltered for hours. We have used 100 ensembles to produce the length distribution and related statistical quantities. For additional parameter values see Table 1 and Table 2.

In the presence of capping proteins, the filament growth dynamics are described by Reactions 2-6 and Reaction 15 (Fig. 1). The capping bound filaments do not add or lose any monomers (i.e., *K*^on^, *K*^off^ = 0) and remain unchanged in length. Here we have ignored the assembly dynamics via pointed end of the F-actin and we shall discuss the effect of pointed end growth later in the study.

**Table 1:**
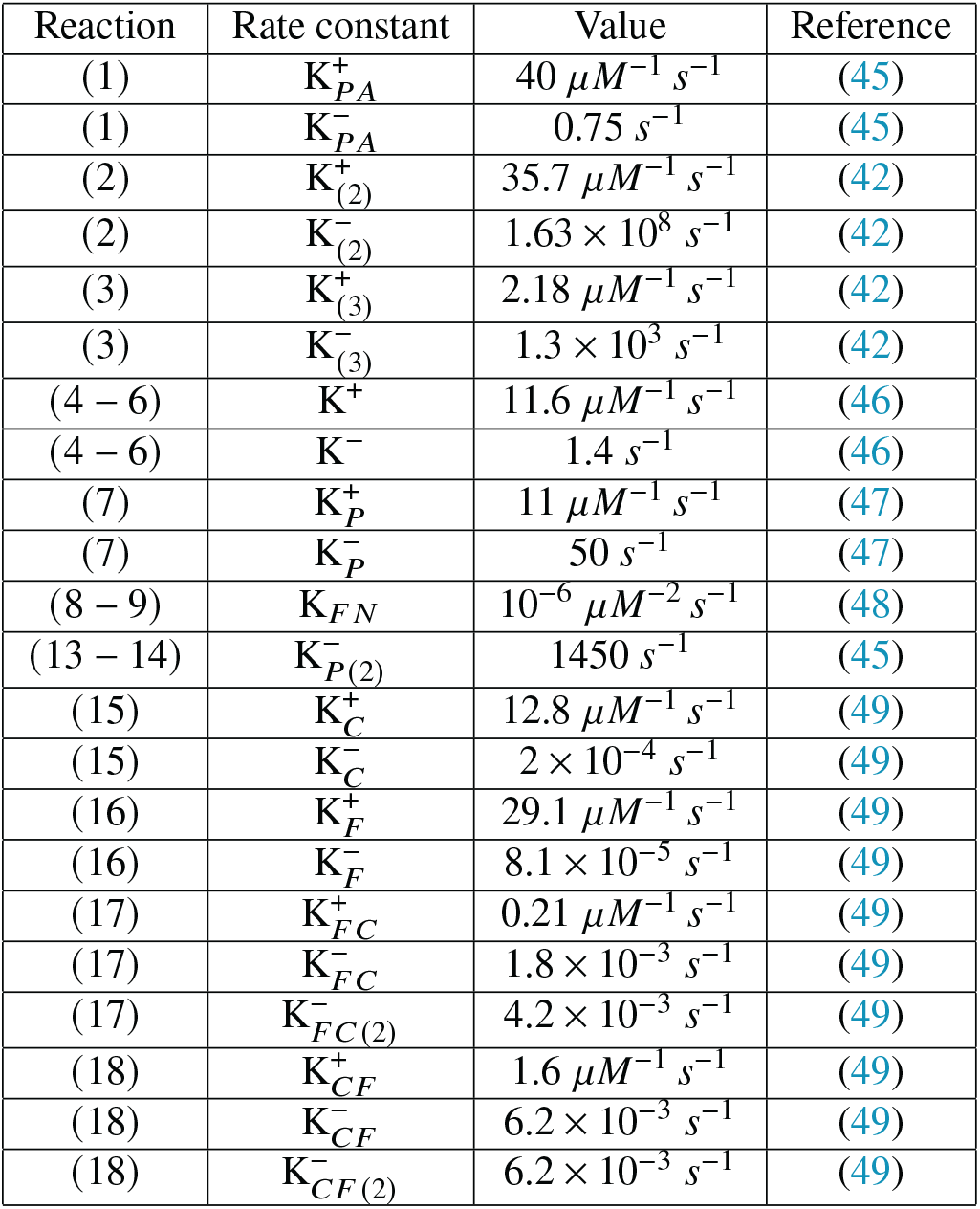
Rate constants used in model simulations

| Reaction | Rate constant | Value | Reference |
| --- | --- | --- | --- |
| (1) | $K_{PA}^+$ | $40 \mu\text{M}^{-1} \text{s}^{-1}$ | (45) |
| (1) | $K_{PA}^-$ | $0.75 \text{s}^{-1}$ | (45) |
| (2) | $K_{(2)}^+$ | $35.7 \mu\text{M}^{-1} \text{s}^{-1}$ | (42) |
| (2) | $K_{(2)}^-$ | $1.63 \times 10^8 \text{s}^{-1}$ | (42) |
| (3) | $K_{(3)}^+$ | $2.18 \mu\text{M}^{-1} \text{s}^{-1}$ | (42) |
| (3) | $K_{(3)}^-$ | $1.3 \times 10^3 \text{s}^{-1}$ | (42) |
| (4 – 6) | $K^+$ | $11.6 \mu\text{M}^{-1} \text{s}^{-1}$ | (46) |
| (4 – 6) | $K^-$ | $1.4 \text{s}^{-1}$ | (46) |
| (7) | $K_P^+$ | $11 \mu\text{M}^{-1} \text{s}^{-1}$ | (47) |
| (7) | $K_P^-$ | $50 \text{s}^{-1}$ | (47) |
| (8 – 9) | $K_{FN}$ | $10^{-6} \mu\text{M}^{-2} \text{s}^{-1}$ | (48) |
| (13 – 14) | $K_{P(2)}^-$ | $1450 \text{s}^{-1}$ | (45) |
| (15) | $K_C^+$ | $12.8 \mu\text{M}^{-1} \text{s}^{-1}$ | (49) |
| (15) | $K_C^-$ | $2 \times 10^{-4} \text{s}^{-1}$ | (49) |
| (16) | $K_F^+$ | $29.1 \mu\text{M}^{-1} \text{s}^{-1}$ | (49) |
| (16) | $K_F^-$ | $8.1 \times 10^{-5} \text{s}^{-1}$ | (49) |
| (17) | $K_{FC}^+$ | $0.21 \mu\text{M}^{-1} \text{s}^{-1}$ | (49) |
| (17) | $K_{FC}^-$ | $1.8 \times 10^{-3} \text{s}^{-1}$ | (49) |
| (17) | $K_{FC(2)}^-$ | $4.2 \times 10^{-3} \text{s}^{-1}$ | (49) |
| (18) | $K_{CF}^+$ | $1.6 \mu\text{M}^{-1} \text{s}^{-1}$ | (49) |
| (18) | $K_{CF}^-$ | $6.2 \times 10^{-3} \text{s}^{-1}$ | (49) |
| (18) | $K_{CF(2)}^-$ | $6.2 \times 10^{-3} \text{s}^{-1}$ | (49) |

**Table 2:** Parameter values

| Figure | Values |
| --- | --- |
| Fig. 2b | $V = 10 \mu\text{m}^3, n_f = 4, \rho_0 = 1 \mu\text{M}$ |
| Fig. 2c-f | $V = 200 \mu\text{m}^3, n_f = 153, \rho_0 = 10 \mu\text{M}$ |
| Fig. 3, 4, 6 | $V = 200 \mu\text{m}^3, \rho_0 = 10 \mu\text{M}, \alpha = 5$ |
| Fig. 5 | $V = 200 \mu\text{m}^3, \rho_0 = 10 \mu\text{M}, \lambda = 6, \gamma = 3$ |

In the presence of profilin and formin, the growth dynamics are given by Reactions 1-14 and Reaction 16 (Fig. 1). For simplicity, we first adopt an effective description where formin-bound filaments assemble with increased rate, given by *K*^on^ = *a:K*^+^ [*A*_1_] and disassemble with the same rate as the free barbed end (Fig. 4). Later, we discuss the effect of profilin and formin-mediated nucleation on filament length dynamics (Fig. 5). To simulate the combined effect of capping, formin and profilin proteins on actin filament growth, we used an effective description (Reactions 2-6 and 15-18 in Fig. 1) in Fig. 6 and a complete description using all the reactions (1-18 in Fig. 1) is presented in Fig S9.

The dimer and profilin-actin dynamics (Reaction 1 and 2 in Fig. 1) are computed using deterministic evolution of concentration of the constituent reactants. We evolve the deterministic equations multiple times to attain equilibrium dimer and profilin-actin concentration between two stochastically chosen reactions *τ* time apart. This treatment of dimer dynamics is similar to previous studies (34). Deterministic evolution of these reactions were adopted to avoid the computationally costly simulation of dimers and profilin-actin dynamics.

The filament length distributions presented in this study are computed from the filament length recorded during the last one minute of the simulation, unless specified otherwise. For most of the study, we consider a representative system with total actin concentration 10 *µ*M and system volume 200 *µ*m^3^. In our study we have not taken spontaneous fragmentation and annealing of filaments into account as these slow processes may not affect the early time (∼30 min) dynamics of filament length at 10 *µ*M actin concentration. It is also not well understood how spontaneous fragmentation and annealing affect actin filament growth dynamics *in vivo* (43), and *in vitro* in the presence of other actin binding proteins. There are conflicting reports from *in vitro* experiments showing exponential (32) and unimodal-like (44) size distribution of actin filaments, observed over a few hours in the absence of any other actin binding proteins..

## RESULTS AND DISCUSSION

### Spontaneous nucleation promotes F-actin length heterogeneity

We first study the length distribution of actin filaments emerging from spontaneous nucleation in a limiting monomer pool without any actin binding proteins, where the filament number and length both evolve in time. The dimers and the trimers have a high dissociation rate, making spontaneous nucleation of actin filaments an inefficient process. In addition, the actin binding protein profilin is known to suppress spontaneous actin nucleation *in vivo*. However, spontaneous filament assembly may play a significant role *in vitro* situations where only actin is present, or when profilin and formin concentrations are relatively low compared to actin.

The growth dynamics for the *i*^th^ filament (i.e., for length *>* 3 monomers) is described by the following chemical master equation

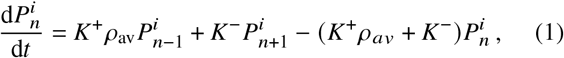

where 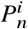 is the probability of the *i*^th^ filament having a length *n* (in monomer units) and *K*^+^ (*K*^−^) is the bare assembly(disassembly) rate. The instantaneous monomer density *ρ*_av_ is given by 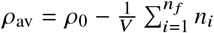 where _0_ is the total actin concentration *V* is the system volume, *n*_i_ is the length of the *i*^*th*^ filament (in monomer numbers), and *n* _*f*_ is the number of actin filaments. The growth of all the filaments can be described by a set of such master equations written according to the polymerization kinetics defined in Fig. 1 and Fig. 2a.

With the parameters listed in Table 2, the nucleation process ends within a few minutes and the mean filament length reaches equilibrium in 10-15 min. Over longer timescales, the filaments can only exchange monomers with other filaments via disassembly and assembly, such that the filament population dwells in a long-lived dynamic state.

In the absence of spontaneous nucleation, when there is a fixed number of filaments simultaneously growing from a limiting monomer pool (i.e., instantaneous nucleation), the resulting length distribution is narrow with a very small coefficient of variation that is maintained over a timescale of 30 min to few hours (Fig. 2b). This early-time length distribution then evolves to become broader, keeping the mean length constant. The distribution eventually becomes exponential over a very long timescale of several days (Fig. 2b). Thus having a large enough G-actin pool can be sufficient to achieve a well regulated length for multiple filaments growing from a shared subunit pool, which is maintained over physiologically relevant timescales of tens of minutes to a few hours. By contrast, the length distribution is significantly broader when the filaments nucleate via spontaneous formation of trimer seeds (Fig. 2c), with all other conditions being the same. The mean length is the same in both cases (Fig. 2d) as it is a property of the pool size and growth rates. This broadness of length distribution originates from the heterogeneity in length in the filament population (Fig. 2e) and not from temporal fluctuations in individual filament length (see Fig. S2).

Emergence of filament length heterogeneity can be understood from the interplay between sequential nucleation of filaments and their subsequent growth. After nucleation, filaments start growing with a growth rate proportional to the instantaneous monomer pool density and all filaments keep growing until the monomer pool density has reached the critical value *K*^−^ / *K*^+^ 0.12 *µ*M. Thus, the filaments that nucleated earlier attain a much larger length compared to the filaments that are nucleated later (see Movie S1). Once the monomer density has reached its critical value, the filaments can only change their length by exchanging monomers between each other. This diffusive process of length rearrangement is very slow for a large actin pool. This leads to the long-lived dynamic state that retains the heterogeneity in filament length. This consequently results in filament length being correlated with their age (Fig. 2f), which is retained for a very long time (Fig. 2f, inset). This age-length correlation is noteworthy because it presents a possible regulatory mechanism where ABPs (such as ADF/cofilin) that interact with filaments in an age-dependent manner (50) (e.g., nucleotide phosphorylation state dependent) can exploit this correlation to build a length-dependent regulation of filament growth. The length heterogeneity decreases (increases) with increasing (decreasing) rate of nucleation, as we have shown by changing dimer production rate (see Fig. S2). Thus, cellular processes that can affect the nucleation rate can regulate the length heterogeneity of the resulting filament population. It is important to note that the timescale of relaxation of filament length distribution (shown in Fig. 2b) is much different in a cell where numerous factors contribute to faster turnover of actin filaments. Spontaneous fragmentation and annealing of actin filaments, observed in *in vitro* systems (32), may also decrease this relaxation timescale. However, the rates of fragmentation and annealing being very small (order of 10^−7^ *µ*M/s) (32), length relaxation via fragmentation and annealing will take much longer than the typical timescales associated with the turnover of actin-based structures (∼30 min - 1 hr).

### F-actin capping increases filament length heterogeneity

We next investigate how the length distribution of actin filaments are regulated by actin binding proteins that inhibit filament growth. Capping proteins (CP) act as F-actin growth inhibitors by binding to F-actin barbed ends and blocking the assembly and disassembly of monomers from that end (Fig. 3a) (51, 52). Here capping proteins are modeled as coarse-grained moieties that associate F-actin barbed ends at a rate 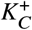 and dissociate at a rate 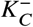 (Reaction 15 in Fig. 1). Capping-bound filaments will stall filament growth (*K*^+^, *K*^−^ → 0) until the capping protein unbinds. We have used Reactions 2-6 and Reaction 15 (Fig. 1) to model the effect of capping proteins, with parameters benchmarked for the well studied capping protein CapZ (Table 1).

**Figure 3:**
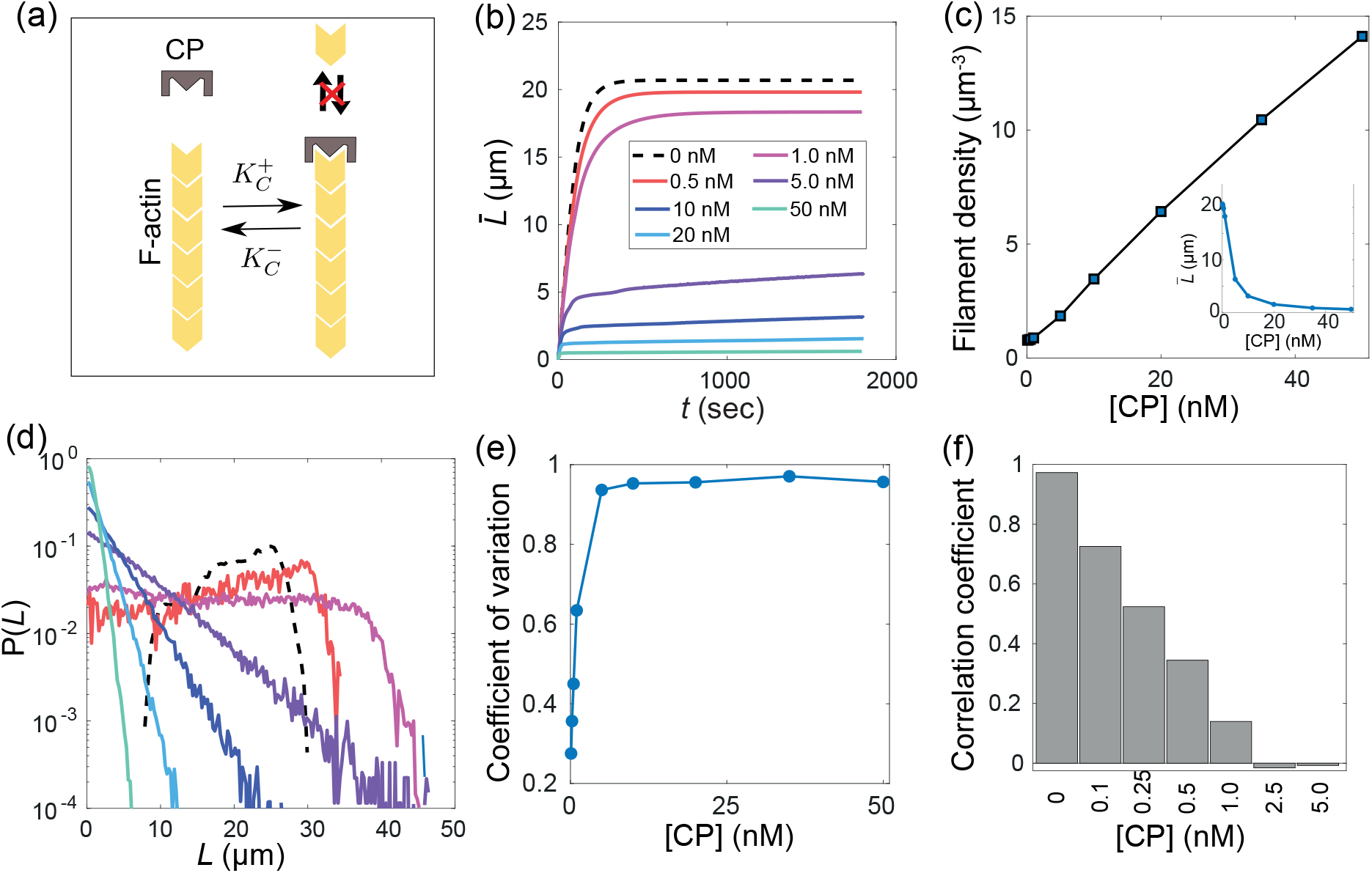
F-actin growth inhibition and length control by capping proteins. (a) Schematic diagram showing capping protein binding to F-actin barbed ends. Capping bound filaments do not grow or shrink in length. (b) Average F-actin length vs time for different concentrations of capping proteins. Increasing capping concentration decreases the mean filament length. (c) Filament number density increases with increasing capping concentration as inhibition of F-actin growth leaves excess amount of actin monomers for spontaneous nucleation. (inset) Mean length of F-actin vs capping protein concentration. The mean length being inversely proportional to capping concentration is a consequence of linear increase in filament numbers with capping concentration. (d) F-actin length distribution for different values of capping protein concentration. The length distribution becomes exponential with increasing capping concentration. The color code is same as shown in panel (b). (e) The coefficient of variation (CV) in filament length vs capping concentration, showing that CV increases with increasing capping concentration and saturates close to one as the distribution becomes exponential. (f) The correlation coefficient between filament age and filament length as a function of capping protein concentration. The filament length and age becomes completely uncorrelated in higher capping concentration. We have used 100 ensembles and collected length for the last one minute to produce the length distribution and related quantities. For additional parameter values see Table 1 and Table 2.

In presence of capping proteins, the mean filament length decreases (Fig. 3b) with increasing capping concentration (*ρ*_*c*_). The mean length 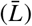 is approximately inversely proportional to *ρ*_*c*_ (Fig. 3c-inset). The reduction in average length of Factin with increasing CapZ concentration cannot be explained without considering the effect of capping proteins on filament nucleation. By inhibiting the growth of the nucleated filaments, the capping proteins induce a slower depletion of the monomer pool and promote nucleation of more filaments (Fig. 3c). The filament density increases approximately linearly with increasing capping concentration (Fig. 3c). This increase in filament abundance decreases the mean length of the filament population. In the absence of spontaneous filament nucleation (i.e., growing a fixed number of filaments), capping proteins can only slow down the growth of the existing filaments. This will lead to the same mean filament length and cannot explain a permanent decrease in average length (Fig. S3).

The filament length distribution loses the unimodal nature in the presence of capping proteins by becoming exponential (Fig. 3d), with the coefficient of variation in length approaching 1 (Fig. 3e). This heterogeneity in length does not arise from exchange of monomers between filaments. Rather, the capping proteins slow down the filament rearrangement dynamics in the diffusive growth regime even further (Fig. S3). The exponential length distribution arises from the interplay between the initial growth of filaments and the dynamics of capping protein binding to the newly created filaments. In the initial period of growth, the filament length grows almost linearly and the waiting time for capping binding to occur has an exponential distribution (as the binding reaction is a Poisson process, independent of filament length). Thus at high enough capping concentration, the filaments captured by the capping proteins will acquire an exponential length distribution. The capping unbinding rate being slow, this nucleation-growth-capture process will deplete the monomer pool and give rise to a long-lived state with large length heterogeneity and an exponential distribution (Movie S2). The process of nucleation-growth-capture does not preserve the information of filament age in their length (Fig. 3f). The capping proteins bind to all free barbed ends with equal probability regardless of the filament age. Hence the difference in filament length emerging previously from the difference in their age, cannot be retained in the presence of capping proteins. The correlation of length with age progressively diminish with increasing capping concentration (Fig. 3f).

In summary, growth inhibition by capping protein enhances heterogeneity in length and we do not observe the long-lived unimodal length distribution that resulted with spontaneous F-actin nucleation (Fig. 2c). Our results are in good agreement with the experimentally measured dynamics of mean filament length (35, 53), length distribution (35) and filament nucleation (54) in the presence of capping proteins. While we neglected actin assembly from the pointed end, pointed-end assembly qualitatively changes actin length distribution in the presence of capping proteins (Fig. S4). In particular, pointed end assembly increases the mean filament length, introduces a peak in the length distribution, while retaining the exponential tail of the distribution.

### Role of Formin on F-actin length control

Assembly of actin filaments in cells is enhanced by growth promoting factors such as formin proteins that bind to the barbed end of the F-actin filament, increasing F-actin polymerization rate up to a few folds (55, 56). Formin proteins not only regulate F-actin organization (30) and actomyosin dynamics (57), but also play important roles in cell motility (58) and cell adhesion (59). While many different types of formins with varied effects on barbed end assembly rate are present in cells, we parametrize our simulations based on mDia1 (49). We model formins as coarse-grained moieties that bind to the barbed end of F-actin at a rate 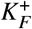 and dissociate at a rate 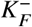 (Fig. 4a), with 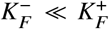. Formin proteins act as growth promoters in the presence of profilin proteins. First, we consider an effective model without explicitly accounting for profilins. In this simplified model, formin-mediated elongation is captured via an enhanced assembly rate *a:K*^+^, with *a: >* 1 (55, 56), allowing us to study the effect of increased elongation rate on filament length distribution. For simplicity, we also neglect the effects of formin-mediated F-actin nucleation. This is a reasonable assumption for some types of formins (e.g., DAAM1, FMNL2 and FMNL3) that have a very low efficiency of nucleating filaments (∼1%) and rather promote barbed end elongation at a higher rate (60–63). We use Reactions 2 - 6 and Reaction 16 (Fig. 1) to model filament growth driven by formin-mediated elongation in the first part of this section (Fig. 4a). We study the role of profilin and formin-mediated nucleation later in this section, where we use Reactions 1 - 14 and Reaction 16 (Fig. 1) to model the filament growth in the presence of both profilin and formin.

**Figure 4:**
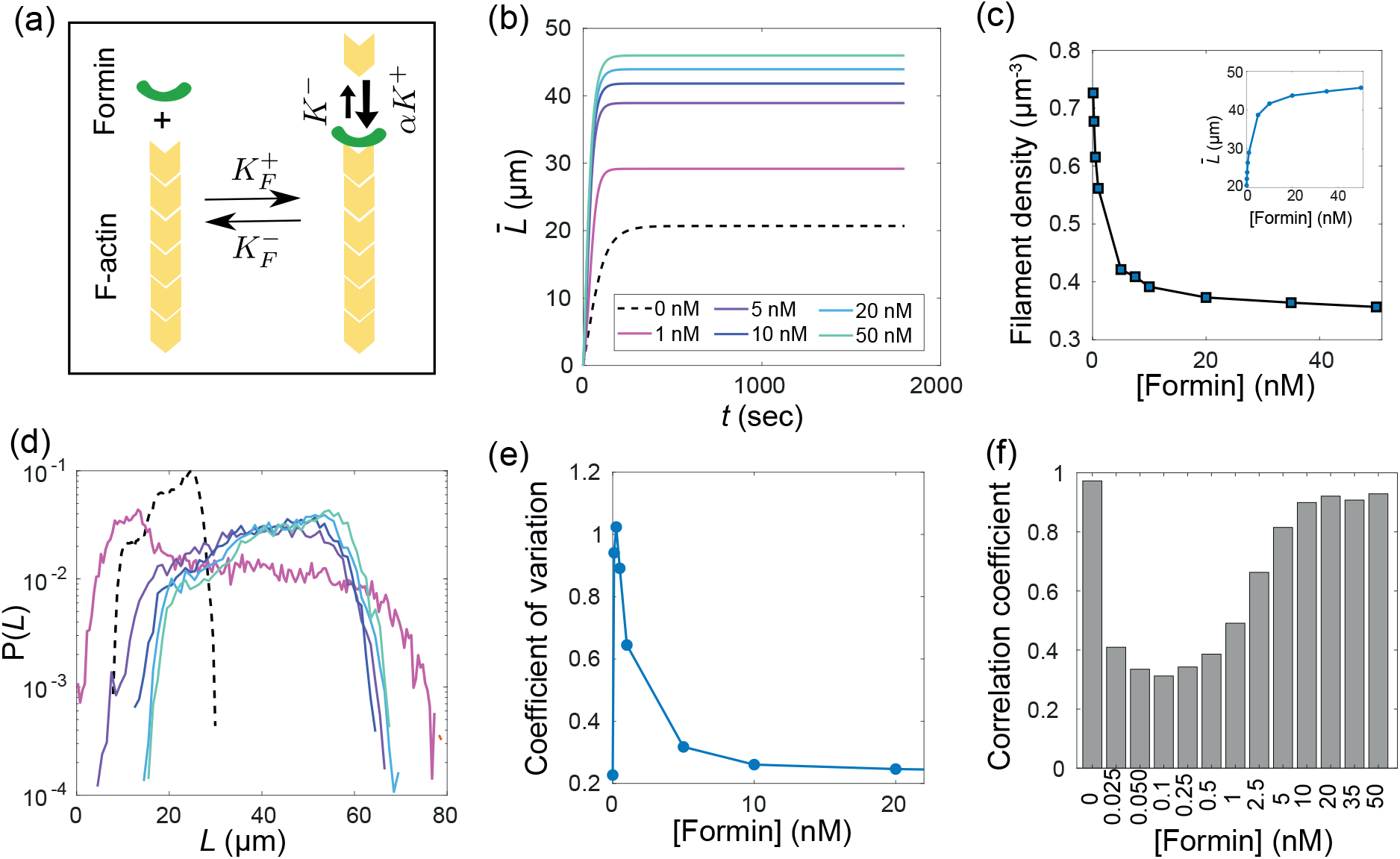
Role of F-actin growth promoters on length control. (a) Schematic diagram of formin binding to F-actin barbed ends, leading to an increase in polymerization rate. (b) The mean filament length increases with increasing formin concentration. (c) The filament number density decreases with increasing formin concentration as promotion of filament growth depletes the monomer pool quicker, leaving less monomers for nucleation. (inset) Increasing filament mean length saturates at higher formin concentration as the filament number density saturates. (d-e) Enhanced polymerization of F-actin by formin retains the unimodality of the filament length distribution but the mean increases. The length heterogeneity is non-monotonic, increasing at lower formin concentration while decreasing later at higher formin concentration. The non-monotonic change in coefficient of variation captures the increased heterogeneity in length at small formin concentration. The color code is same as shown in panel-b. (f) The correlation between filament age and length is non-monotonic, with a decrease at an intermediate range of formin concentrations. We have used 100 ensembles and collected length for the last one minute to produce the length distribution and related quantities. For additional parameter values see Table. 1 and Table. 2.

#### Role of Formin-mediated elongation

We find that the mean length (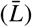) of growing actin filaments increase with increasing formin concentration (Fig. 4b), while eventually saturating at higher values of formin concentration (Fig. 4c-inset). This saturation of mean length occurs as nucleated filaments remain formin-bound most of the time at high formin concentration, making the effective actin assembly rate ∼ *a:K*^+^. Presence of formins can change the critical concentration for actin growth in a concentration-dependent manner (see Fig. S5). Depending on the change in elongation rate mediated by different types of formin proteins, the critical concentration may increase or decrease compared to the critical concentration in the presence of only actin. However, these estimates of critical concentration are based on our effective description may not accurately capture the critical concentration in real experimental systems where non-equilibrium processes regulate various aspects of actin assembly. The enhanced growth of the formin-bound filaments leads to an accelerated depletion of the monomer pool, thereby suppressing spontaneous nucleation. Filament density decreases with increasing formin concentration, approximately scaling inversely (Fig. 4c).

With increasing formin concentration, filament length distribution shifts to a higher mean but remains unimodal and qualitatively similar to the case without formin (Fig. 4d). At high enough formin concentration, formins strongly promote growth of the filaments, preserving the unimodality of length distribution without increasing length heterogeneity (Fig. 4d,e and Movie S3). However, the coefficient of variation in filament length changes non-monotonically with formin concentration, indicating higher heterogeneity at lower formin concentration (Fig. 4e). This increased heterogeneity in length is caused by the competition between formin-bound and free filaments in absence of enough formins to bind all the filaments that are being nucleated (see Fig. S5). This non-monotonicity is also present in the correlation between filament length and age, with a loss of correlation at small formin concentration, while regaining the correlation back at higher concentration of formins (Fig. 4f). The similarity in length-age correlation at high and low formin concentrations arises from the fact that at high formin concentration the actin growth can be effectively represented as growth of only F-actin with an enhanced assembly rate (∼ *a:K*^+^).

#### Role of profilin and Formin-mediated nucleation

Formin is known to increase the assembly rate of F-actin in the presence of profilin proteins that bind to actin monomers to create a profilin-actin (PA) pool (64). PA selectively takes part in the assembly of formin-bound filaments with higher assembly rate. In the above, we did not explicitly consider profilin dynamics and implicitly assumed that formin-bound filaments incorporate PA to increase their assembly rates. We also assumed that spontaneous nucleation can proceed in the presence of profilin. These assumptions are reasonable if the amount of actin monomers is much larger than profilin-actin amount in the pool. Actin-profilin ratios in cells are typically larger than 2:1 (45), leaving enough free monomers for spontaneous nucleation in the beginning. However, with the growth of formin-bound filaments the profilin-actin subunits quickly become dominant over free actin monomers.

To study the role of profilin-actin on F-actin length control, we explicitly modeled profilins and their interaction with actin (Fig. 1). In addition, we modeled irreversible dimer formation (from either two G-actin or one G-actin and one profilin-actin) by formins at a rate *K*_FN_ (see Supplemental Methods for details) to study the role of formin-mediated nucleation on F-actin length distribution. We considered three different profilin concentrations of 2 *µ*M, 5 *µ*M and 10 *µ*M, corresponding to the physiologically relevant actin-profilin ratios 5:1, 2:1, and 1:1, respectively. We find that the mean filament length shows a non-monotonic dependence on formin concentration, results from a competition between spontaneous nucleation and formin-mediated F-actin nucleation (Fig. 5a). Filament length is maximum at an intermediate formin concentration, when formin-mediated nucleation is not prevalent but most of the filaments are formin-bound and grow to be large. At a fixed formin concentration, mean length increases with increasing profilin concentrations due to less spontaneous nucleation and faster elongation rate of formin-bound filaments (Fig. 5a,b). While the length distributions are more spread-out at higher actin-profilin ratios (Fig. 5b), length heterogeneity remains almost invariant for changing formin concentration and actin-profilin ratio. At very low formin concentration, length heterogeneity is high due to competing subpopulations of filaments with formin-bound and free barbed ends (Fig. 5c and Fig. S6). At high profilin concentrations (10 *µ*M with actin-profilin concentration ratio 1:1), the mean length decreases with increasing formin concentration as more filaments are created via formin-mediated nucleation (Fig. 5d). As reported in previous studies (48), a stronger formin-mediated nucleation leads to an increase in filament density (Fig. 5e) and a reduction in mean filament length (Fig. 5f) with increasing formin concentration. At lower formin concentrations and smaller rate of formin-mediated nucleation, *K*_FN_, mean filament length increases due to reduced nucleation of filaments (see Fig. S7 & S8). This is similar to the case where formins act effectively as growth promoters (Fig. 4).

**Figure 5:**
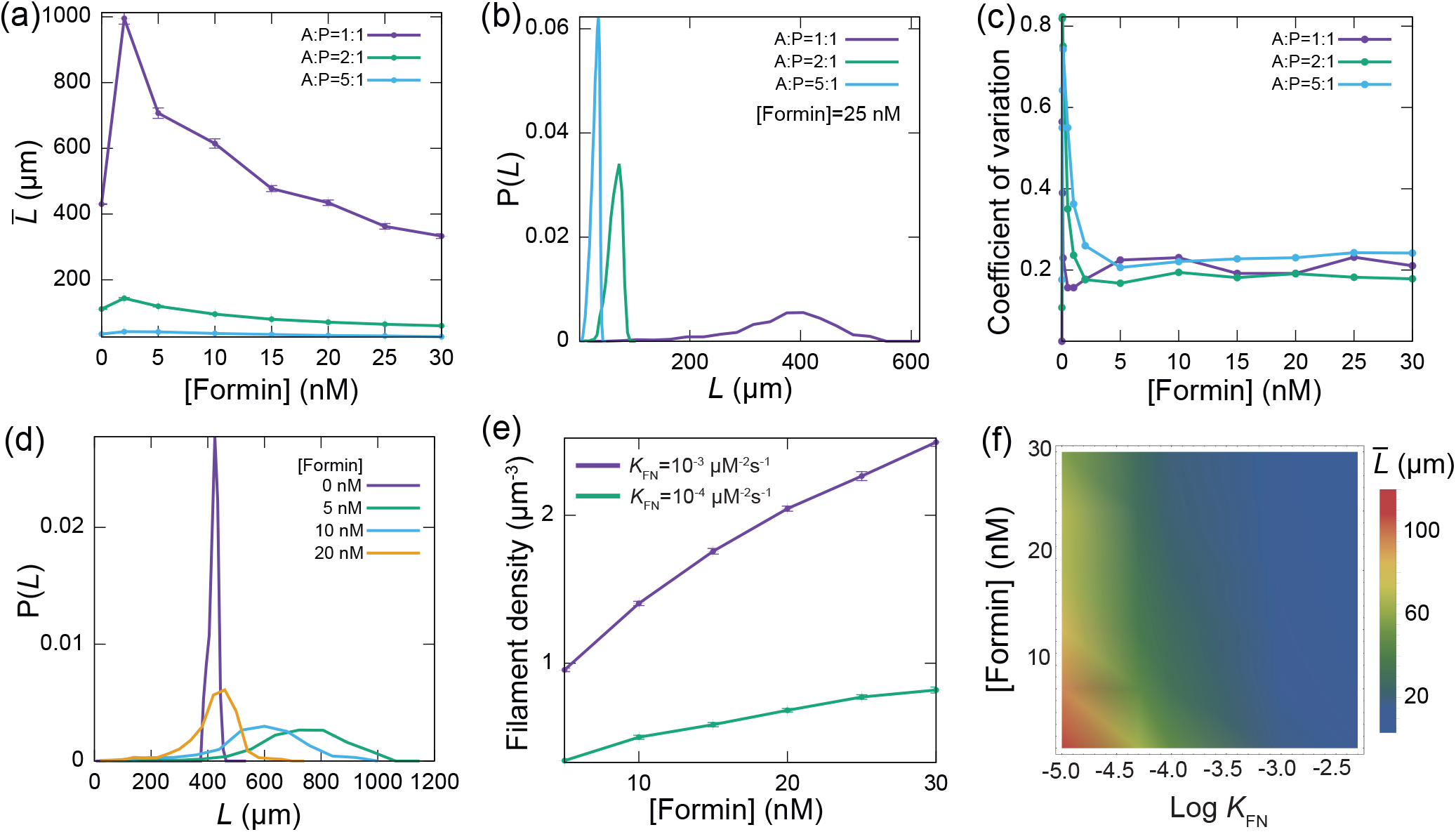
Effects of actin sequestration by profilin and formin-mediated nucleation on actin filament growth. (a) Mean length of filaments as a function of formin concentration for three different actin-profilin concentration ratios. (b) Length distribution of actin filaments for different actin-profilin concentration ratios at a fixed formin concentration. (c) Length heterogeneity, measured by coefficient of variation, changes non-monotonically with formin concentration with a higher length heterogeneity at very small formin concentration. Length heterogeneity at larger formin concentrations is small and becomes independent of formin or profilin concentrations. (d) Length distribution of actin filaments for different formin concentrations at the actin-profilin ratio 1 : 1 (i.e., actin and profilin concentrations are same and equal to 10 *µ*M). (e) Filament density increases with formin concentration in the presence of strong formin-mediated nucleation. (f) Dependence of mean actin filament length as a function of formin-mediated dimerization rate (*K* _*N*_) and formin concentration. The heatmap shows that mean length decreases with increasing *K* _*N*_ and formin concentration. The actin-profilin concentration ratio is chosen to be 2:1 in panels (e) and (f). For additional parameter values see Tables 1 and 2.

Overall, our results predict the effects of filament elongation and nucleation by formins on the emergent length distribution of actin filaments. The predicted Filament length dynamics in the presence of formin-mediated nucleation (Fig. 5) are in good agreement with the recently reported role of formin in limiting actin filament length in an *in vitro* assay (48). The results from our effective description of formins as growth promoters (Fig. 4) may be relevant *in vivo* where formin-mediated nucleation often requires other co-factors or nucleation promoting factors. In the absence of such cofactors, formins only play an essential role in barbed-end elongation. For mDia1 the co-factor required for nucleation is adenomateous polyposis coli (APC) and without APC, mDia1 hardly nucleates new filaments in the presence of profilin (65).

### Competition between Formin and Capping proteins results in bimodal F-actin length distribution

In the previous sections, we studied how F-actin growth inhibitors and growth promoters maintain filament length heterogeneity and retain unimodality of length distribution. Here we study their combined effect on filament growth and length control. To model the interactions between formin and capping proteins, we adopt an effective description for formin-driven F-actin elongation as before (Reactions 2 − 6, 15 − 18 in Fig. 1), without explicitly accounting for profilin and formin-mediated nucleation. Formin and capping proteins both compete for the F-actin barbed end and this competition was previously thought to be exclusive (66). Recent studies (49, 67) have uncovered the interaction between free formins (F) and capping bound to filaments (BC) and vice-versa. When bound to a barbed end, formin and capping proteins can form a ternary complex, referred to as BFC or BCF depending on the order of the complex formation (Fig. 6a). For instance, capping protein binding to a formin-bound barbed end forms a BFC complex, while formin binding to a capping-bound barbed end forms a BCF complex. These complexes can disassemble in two different ways leaving the barbed end either formin-bound or capping-bound (Fig. 6a). The disassembly rate of this complex is larger than the very small disassembly rate of capping and formin proteins from the barbed end (Table 1). Thus this interaction enables actin filaments to switch from a capping-bound non-growing state to a formin-bound fast growing state.

**Figure 6:**
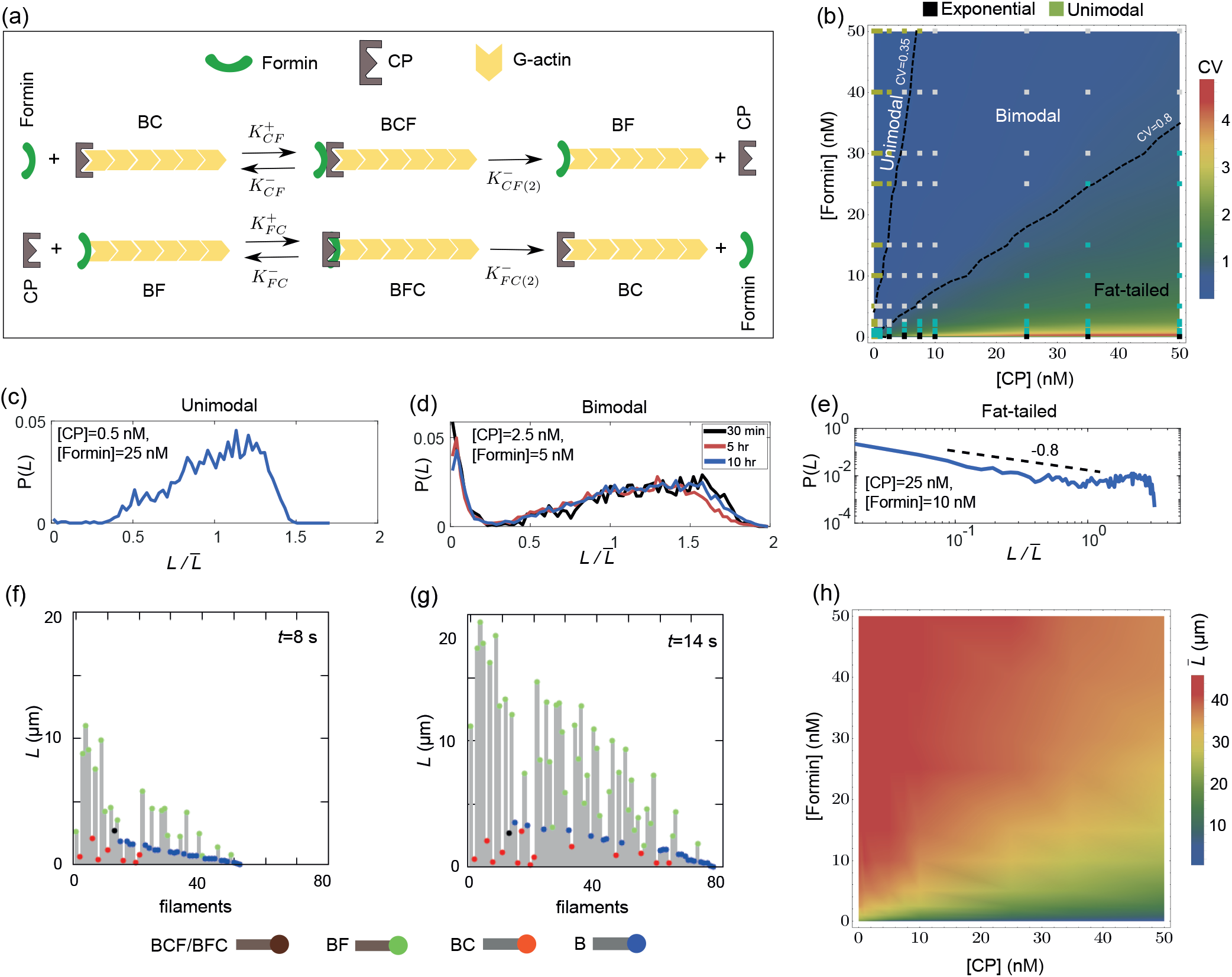
Emergence of bimodal F-actin length distribution by competition between formin and capping proteins. (a) Schematic diagram of the interaction between actin, formin and capping proteins. The capping-formin complex may form in two different ways, either by formin binding to a capping-bound barbed end (BC) or vice-versa, with different rates of association and dissociation. (b) A state-diagram showing the different F-actin length distributions resulting at different values of formin and capping protein concentration. Four qualitatively different distributions emerge: Unimodal, exponential, bimodal and fat-tailed distribution. The fat-tailed length distribution has very broad power-law tails and a large heterogeneity in length. (c-e) Representative plots for the three types of length distributions (left to right): (c) unimodal, (d) bimodal and (e) fat-tailed distributions computed at the indicated values of formin and capping protein concentrations. The bimodal length distribution (at [Capping]=2.5 nM and [Formin]= 5 nM) remains almost unchanged over a long time period spanning minutes to tens of hours. (f-g) Segregation of filaments during early growth period into subpopulations of filaments of large and small length by formin and capping, respectively. Shown here are the filament length distributions at (f) *t* = 8s and (g) *t* = 14s. The parameters used are [Capping]=4 nM and [Formin]= 5 nM. (h) Mean filament length as a function of formin and capping protein concentrations. Increasing formin (capping) concentration increases (decreases) the mean length of the filament population. Additional parameter values are given in Table 1 and Table 2.

By varying the amount of formin and capping proteins, we find four distinct types of F-actin length distributions that emerge via the interaction of capping and formin proteins with the F-actin filaments (Fig. 6b). When formin concentration is considerably higher than the capping protein concentration, the resulting length distribution is unimodal (Fig. 6c). Here, the formin proteins outcompete the effect of capping proteins and retain unimodality of length distribution. We find the characteristic exponential length distribution emerging from capping-induced inhibition of F-actin growth, but this effect fades away in the presence of even small amount of formin (Fig. 6b). In the region where concentration of capping proteins is much higher than formin proteins we find that the filament population has a high amount of heterogeneity in length and the coefficient of variation is large (*CV >* 1) (Fig. 6b and Fig. 6e). We refer to these length distributions with powerlaw tails followed by an eventual exponential decay (Fig. 6e, Fig. S9) as fat-tailed distributions. This broadness in length distribution appears where amount of capping is higher than formin but it is noteworthy that capping proteins alone cannot produce such high amount of heterogeneity in length and the competition between capping and formin proteins plays a role in promoting this large heterogeneity in length.

When formin and capping protein concentrations are comparable, we see the emergence of a bimodal length distribution, where the filament population has two clearly segregated subpopulations of small and large filaments (Fig. 6b and Fig. 6d).

The bimodal length distribution is long-lived, being stable without any significant changes over a timescale of tens of minutes to tens of hours (Fig. 6d). The origin of the bimodality lies in the early time segregation of filaments either by growth promoters in the subpopulation of large filaments or by growth inhibitors in the subpopulation of small filaments (Fig. 6f-g and Movie S4). These segregated subpopulations are stable over a timescale of many hours, as the monomer pool reaches critical concentration when it is in equilibrium with the filaments. After this initial period of growth, formin and capping proteins cannot significantly alter the filament length distribution, as in this long-lived diffusive growth regime a small filament cannot grow fast to become large even if it is formin-bound. In addition to modulating the nature of length distributions, the mean length of the filament population can also be tuned by changing formin and capping protein concentrations (Fig. 6h). Increase in formin concentration and decrease in capping protein concentration increases the mean filament length as expected from the results of the previous sections.

We tested the validity of our assumption of neglecting the role of profilin and formin-mediated nucleation by explicitly modeling these effects. Our results show that the qualitative nature of the filament length distribution and the phase diagram do not change in the presence of profilin and formin-mediated nucleation, such that we can obtain both bimodal and fat-tailed length distributions (see Fig. S6 & S10).

## CONCLUSIONS

In this article, we developed a stochastic model of actin filament nucleation and assembly from a shared limiting pool of monomers and binding partners. Using this model, we delineate the conditions under which actin filaments of variable lengths can be built and maintained. Although we ignore many important actin regulators relevant *in vivo*, our model provides valuable insights into how actin length can be dynamically controlled by tuning concentrations of Factin nucleators, growth promoters and growth inhibitors. Our model can thus be better realized in an *in vitro* setup, where the concentrations of actin and its binding partners can be precisely controlled. Our computational model can be extended further by incorporating F-actin fragmentation and annealing, which have shown to affect F-actin length distribution over longer timescales *in vitro* (32). While the role of spontaneous F-actin fragmentation and annealing remains unclear *in vivo* (43), models can shed light into how these slow processes may regulate filament length distribution in the presence of growth promoters and growth inhibitors.

We show that spontaneous nucleation can induce significant length heterogeneity in the filament population. This heterogeneity in length is an emergent property of spontaneous nucleation that cannot result from the growth of a fixed number of filaments from specialized nucleators (26, 27). In addition, spontaneous nucleation results in a strong age-length correlation in the filament population. A recent study by Rosenbloom *et al* (68) reports much slower formation and degradation of actin dimers than was previously reported by Sept and McCammon (42). With slower nucleation rates, we find that the mean length of actin filaments increases (resulting from smaller filament density), while the qualitative features of the filament length distribution remained unchanged (Fig S11).

Filament length heterogeneity and distribution can be controlled by tuning the interaction between F-actin and its growth promoters and inhibitors. We find that strong growth inhibition leads to increased length heterogeneity within physiologically relevant timescales. In this case, length-independent growth inhibition by capping proteins reduces the age-length correlation in the filament population and a complete loss of correlation is seen at higher concentration of capping proteins. Our results agree well with *in vitro* studies reporting exponential length distribution and decreasing mean filament length with increasing capping concentration (35). Enhanced F-actin elongation by formin reveals a non-monotonic behaviour in filament length distribution with increasing formin concentration. The heterogeneity in length increases for small concentration of formin due to a competition between formin-bound and formin-free filaments. This heterogeneity goes away at higher concentration of formins when there are enough formin to bind all the filaments. Thus, a strong enhancement of growth rate does not result in any significant change in length heterogeneity and retains the unimodality of length distribution.

Motivated by recent *in vitro* studies reporting the interactions between formin and capping proteins (49, 67), we studied actin filament growth in the presence of formin and capping proteins. We show that formin and capping protein concentrations can be tuned to regulate the heterogeneity in length in the filament population. We find a bimodal length distribution in a regime where the concentration of formin and capping proteins are comparable such that their competition is enhanced. Bimodal length distribution may originate from an underlying bistability due to autocatalytic growth (27), as observed *in vitro* for microtubules growing in the presence of Kip3 motors (69). However, the bimodal length distribution we report here does not originate from any underlying bistability in the filament growth dynamics. Here bimodality is an emergent collective effect that arises from competition between actin growth promoters and growth inhibitors. This mechanism for maintaining subpopulations of short and long actin filaments may play an important role in assembling distinct actin network organizations with variable lengths (70, 71). The filaments observed *in vivo* are often much smaller than the range of filament lengths reported in our simulations (Fig. 6h). This is because actin density inside the cell are much higher than what we considered, and numerous other actin regulators in the cell may further reduce filament lengths by enhanced nucleation (e.g. by Arp-2/3) or via F-actin severing (e.g. via ADF/cofilin).

We show that explicit modeling of profilin in physiologically relevant concentrations does not alter the qualitative results but changes the overall length of the formin-bound subpopulation of filaments (see Fig. S6). By accounting for formin-mediated filament nucleation, we find that the dependence of filament length on formin concentration is altered, depending on the formin nucleation efficiency. But the qualitative features of the filament length distributions do not change in different growth regimes. Interestingly, while formin retains length heterogeneity and capping proteins increase it, profilins are able to reduce filament length heterogeneity by limiting nucleation (see Fig. S12). Our model predicts a monotonic increase in actin elongation rate with increasing profilin concentration (Fig. S12), which is in good agreement with experimental data for profilin concentrations lower than that of actin (72). At even higher profilin concentrations, actin elongation rate is found to decrease with profilin concentration (72), which is not captured by our current model since we do not consider the interaction between free profilins and formin.

To the best of our knowledge, bimodality in actin length distribution has not yet been experimentally observed in the presence of formin and capping. The recent studies on capping-formin interaction (49, 67) are done in flow channels where the monomer pool may not be conserved and data on filament length distribution is not reported. Interestingly, Zigmond *et al* (66) found unimodal length distribution in the presence of 0.5 *µ*M actin, 10 nM capping and 200 nM formin, which is in good agreement with our prediction that the length distribution is unimodal when formin concentration is much larger than that of capping proteins (Fig.6b). Two kinetically distinct (different turnover rates) actin filament subpopulations regulated by arp-2 / 3 and formin were reported in cortical actin network (73, 74), but the average lengths of these subpopulations was found to be independent of the concentration of the regulators in the simulations (73). The formin-nucleated filaments were reported to be longer and was found to play an essential role in determining mechanical properties of the cortical network in comparison with the shorter filaments (73). This highlights the significant role that filament length heterogeneity may play in regulating the mechanical properties of actin networks, opening doors to future studies on this topic.

## CODE AVAILABILITY

Custom simulation codes used to generate each figure are available at https://github.com/BanerjeeLab/Actin_Length_Simulations.

## AUTHOR CONTRIBUTIONS

D.S.B. and S.B. designed and developed the study. D.S.B. carried out simulations and analyzed the data. D.S.B. and S.B. wrote the article.

## ACKNOWLEDGMENTS

SB acknowledges funding from the Human Frontier Science Program (HFSP RGY0073/2018), and the National Science Foundation (NSF MCB-2203601). SB and DSB thank Darius Köster and Markus Deserno for useful inputs. DSB thanks Amit Das and Suman G Das for helpful discussions.

## SUPPORTING MATERIAL

An online supplement to this article can be found by visiting the preprint website.

## Movie title and legends

**Movie S1**. Actin filament growth by spontaneous nucleation. The y-axis denotes filament length in *µ*m and the filaments are ordered according to their age, i.e., the right-most filament is the youngest. The correlation in filament age and length can be clearly seen to be emerging from spontaneous nucleation of filaments. The blue dots represent free barbed end. Parameter values used are same as in Fig. 2c

**Movie S2**. Actin filament growth by spontaneous nucleation in the presence of capping proteins. The y-axis denotes filament length in *µ*m and the filaments are ordered according to their age, i.e., the right-most filament is the youngest. The emergence of heterogeneity, due to capping proteins blocking the growth of newly nucleated filaments, can be seen. The blue and red dots represent the free barbed ends and capping bound barbed ends respectively. Parameter values used are same as in Fig. 3 with capping concentration of 2 nM.

**Movie S3**. Actin filament growth by spontaneous nucleation in the presence of formin proteins. The y-axis denotes filament length in *µ*m, and the filaments are ordered according to their age, i.e., the right-most filament is the youngest. The length heterogeneity and the correlation in filament age and length can be seen to be very similar with the case of spontaneous growth without any ABPs. The blue and green dots represent the free barbed end and formin bound barbed ends, respectively. Parameter values used are same as in Fig. 5 with profilin and formin concentration of 1 *µ*M and 0.5 nM respectively. The formin mediated nucleation rate was taken to be *K*_F_ _*N*_ = 10^−5^ *µM*^−2^*s*^−1^.

**Movie S4**. Actin filament growth by spontaneous nucleation, in the presence of formin and capping proteins. The y-axis denotes the filament length in *µ*m and the filaments are ordered according to their age, i.e., the right-most filament is the youngest. The emergence of two subpopulations of short and long filaments can be seen. The subpopulations arise due to the capture of filaments by formin (grows long) and capping (remains small) during the early period of their growth. The blue, green and red dots represent the free barbed end, formin-bound barbed ends, capping-bound or capping-formin complex bound barbed ends, respectively. Parameter values used are same as in Fig. 5 with profilin, capping and formin concentration of 1 *µ*M, 8 nM and 5 nM respectively. The formin mediated nucleation rate was taken to be *K* _*N*_ = 10^−5^ *µM*^−2^*s*^−1^.

