## Supporting Material for "Emergence and maintenance of variable-length actin filaments in a limiting pool of building blocks"

### Supplemental Methods

#### Stochastic filament growth model

We use the Gillespie algorithm [1] to simulate the stochastic assembly/disassembly reactions except dimer formation and degradation which was computed by deterministically updating the reactant numbers (discussed later). The classic Gillespie algorithm uses two random variables at any time $t$, drawn from an uniform distribution ($r_1, r_2 \in \mathscr{U}(0,1)$), and the instantaneous propensities for all of the possible reactions to update the system in time according to the defined growth law. The propensities of the relevant growth reactions, i.e., the nucleation, assembly and disassembly rates of the $i^{th}$ filament (i.e., length larger than 3 in monomer units) are given by $K^{\text{on}}_{\text{Nuc}}$, $K^{\text{off}}_{\text{Nuc}}$, $K^{\text{on}}_{i}$ and $K^{\text{off}}_{i}$, respectively. For actin filaments growing in a limiting monomer pool, these propensities are functions of the instantaneous cytoplasmic monomer concentration ($\rho_{\text{av}}$ or $[A_1]$ as will be used here) and bare growth rates (described in main-text Table 1) which depend on the state of the barbed end (i.e., if it is free or ABP bound). For nucleation (i.e., formation of a trimer) the rates are

$$
\begin{aligned}
K^{\text{on}}_{\text{Nuc}} &= K^{+}_{(3)}[A_2][A_1] \ , \\
K^{\text{off}}_{\text{Nuc}} &= K^{-}_{(3)}[A_3] \ , \qquad (1)
\end{aligned}
$$

where $[A_2]$ and $[A_3]$ are the concentration of dimer and trimer. Filaments with free barbed ends assemble and disassemble with rates given by

$$
\begin{aligned}
K^{\text{on}}_{i} &= K^{+}[A_1] \ , \\
K^{\text{off}}_{i} &= K^{-} \ , \qquad (2)
\end{aligned}
$$

Formin bound filaments assemble with increased rate, given by $K^{\text{on}} = \alpha K^{+}[A_1]$ and disassemble with rate same as the free barbed end. The capping bound filaments do not add or lose any monomers (i.e., $K^{\text{on}}, K^{\text{off}} = 0$) and remain unchanged in length.

The Gillespie algorithm computes the time for the next reaction at $t + \tau$ given the current state of the system (i.e., the propensities for all reactions) at time $t$ where $\tau$ is given by-

$$
\tau = \frac{1}{\sum_{i=1}^{C} \mathscr{R}_i} \log\left(\frac{1}{r_1}\right) \ , \qquad (3)
$$

where $\mathscr{R}_i$ is the propensity of $i^{th}$ reaction and $C$ is the total number of all possible reactions. The second random variable $r_2$ is used to select the particular reaction ($j^{th}$ reaction) that will occur at $t+\tau$ time such that

$$\frac{\sum_{i=1}^{j-1}\mathscr{R}_i}{\sum_{i=1}^{C}\mathscr{R}_i} \leq r_2 < \frac{\sum_{i=1}^{j}\mathscr{R}_i}{\sum_{i=1}^{C}\mathscr{R}_i} \; . \tag{4}$$

The condition for the first reaction ($j=1$) is $0 \leq r_2 < \frac{\mathscr{R}_1}{\sum_{i=1}^{C}\mathscr{R}_i}$. The two steps defined by Eq. 3 and Eq. 4 are used recursively to compute the growth dynamics in time.

The dimer formation and decay are computed using deterministic evolution of dimer concentration $[A_2]$ with rates

$$K_{\text{dim}}^{\text{on}} = K_{(2)}^{+}[A_1]^2 \; ,$$
$$K_{\text{dim}}^{\text{off}} = K_{(2)}^{-}[A_2] \; . \tag{5}$$

We evolve the dimer equation multiple times to attain equilibrium dimer concentration between two stochastically chosen reactions $\tau$ time apart. This treatment of dimer dynamics is similar to previous studies [2]. Deterministic evolution of the dimer formation and decay was adopted to avoid the computationally costly *fast* dissociation of dimers.

#### Sequestration by profilin and formin-mediated nucleation

The schematic in main text (Fig. 1) describes all the reactions involved in actin growth in the presence of capping, formin and profilin proteins, as well as formin-mediated actin nucleation. Below we provide additional details of each of these components.

**Profilin dynamics.** In the main text we have discussed the effects of formin-dependent elongation on filament length heterogeneity without explicitly considering profilin proteins. In reality, formins need profilin-actin monomers for the rapid elongation of F-actins by formins [3]. Profilin proteins bind with acin monomers and sequester them by preventing the formation of dimers (hence disrupt the spontaneous nucleation process) [4]. We implicitly assumed that profilins are present in moderate amounts such that they do not hinder spontaneous filament nucleation yet provide a profilin-actin monomer pool for formins to incorporate actin monomers into filaments with enhanced rate. To verify our assumption we studied actin growth in presence of actin, formin and profilin. To model the interactions of profilin with monomeric actin, actin filament with free barbed end and formin-bound actin filament we have followed the recent study by Funk *et al* [5]. We do not take the interaction of free profilin with barbed end (free or formin bound) as probability of such reactions are quite small in physiological conditions [5]. Consequently, the slowdown of filament elongation at very high profilin concentration (profilin concentration $2-3$ times higher than actin concentration) cannot be captured by our model.

**Formin-mediated nucleation.** In the results presented in the main text, we have assumed that formin-mediated nucleations are negligible as we have considered Diaphanous-related formins like DAAM1 or mDia1 which are very inefficient in nucleating filaments [6]. For the sake of completeness, we have studied the effect of formin-mediated nucleation in the presence of actin, profilin and capping proteins. Following a recent study by Zweifel *et al* [7] we have modeled formin-mediated nucleation of actin filaments via formin-meadiated formation of actin-actin or actin-profilin-actin dimers which irreversibly grow to be formin-bound filaments (Supplementary schematic). In Zweifel *et al*, a processive formin in yeast, Bni1p (more in elongation efficiency and less in nucleation efficiency compared to Cdc12p) was fitted with a dimer nucleation rate $K_{FN} = 35 \times 10^{-6} \mu M^{-2} s^{-1}$. We have explored a range of $K_{FN} = 1 - 50 \times 10^{-6} \mu M^{-2} s^{-1}$ to show that a low nucleation rate and physiologically relevant profilin amount ([Actin]:[Profilin]$\leq 2:1$) do not alter the qualitative nature of length heterogeneity. However, the behaviour of mean actin filament length and number of nucleated filaments changes if moderate formin mediated nucleation is considered. Our findings are consistent with the experimentally observed behaviour reported in Zweifel *et al*.

**Description of chemical reactions.** Below we describe the chemical reactions illustrated in the Supplementary schematic.

Reaction $1-2$: Interaction of profilin with monomeric actin was modelled using a deterministic description to reduce the computational cost.

Reaction $3-5$: Actin trimer and tetramer dynamics was modeled to facilitate a dynamic creation and destruction of filaments ($A_n$ where $n > 4$). We allowed formin and capping binding for filaments of length 5 and more excluding the tetramers.

Reaction $6-7$: Filaments($A_n$ where $n > 4$) can grow by incorporating G-actin ($A_1$) or profilin-actin ($PA_1$)

Reaction $8-12$: We model formin-mediated nucleation in a similar way as presented in Zweifel *et al* [7], where a formin-bound dimer ($FA_2$) can form either from two G-actin subunits or one G-actin and one profilin-actin subunit. We have taken the assembly rate ($K_{FN}$) for both the reactions to be the same. The dimer then can irreversibly grow to become a formin-bound trimer

($FA_3$) and a formin-bound tetramer ($FA_4$), which can grow to be a new formin-bound filament ($FA_n$) and take part in all the reactions allowed for filaments. It is important to note that despite formin-mediated nucleation being irreversible upto tetramer formation the newly nucleated formin bound filament may get disassembled by the dissociation of formin (leaving the filament as $A_n$) and subsequent monomer dissociations. All the binding reactions, except the dimer formation, in box-e happens with rate $\lambda K_P^+$, which is the rate of incorporating profilin-actin monomers in formin bound filaments.

Reaction $13-14$: Profilin-actin monomers get incorporated in filaments with free barbed ends at a rate $K_P^+$ and the same incorporation happens at an enhanced rate $\lambda K_P^+$ when the filament is formin-bound. We model the profilin unbinding event after profilin-actin binding explicitly (with a rate $K_{P(2)}^-$ for free barbed end otherwise with rate $\gamma K_{P(2)}^-$ when the filament is formin bound) as this step was determined to be rate limiting factor in actin filament growth [5]. The profilin-actin monomer can unbind too from the filament with a rate $K_P^-$. Though free profilin can bind to the barbed end, the small binding rate (compared to profilin-actin binding rate) and the lack of free profilin would make this binding event unlikely [5].

Reaction $15-18$: In our model, capping and formin can bind to barbed end that are not profilin bound. This way profilin unbinding step also limits how fast capping proteins or formin can bind to the barbed end. The formation of the "decision complex" is modeled by following the findings reported in Shekhar *et al* [8] and has been discussed in details in the main text.

[7] M. E. Zweifel, L. A. Sherer, B. Mahanta, and N. Courtemanche, Biophysical Journal (2021).

[8] S. Shekhar, M. Kerleau, S. Kühn, J. Pernier, G. Romet-Lemonne, A. Jégou, and M.-F. Carlier, Nature communications **6**, 1 (2015).

[9] I. Fujiwara, D. Vavylonis, and T. D. Pollard, Proceedings of the National Academy of Sciences **104**, 8827 (2007).

[10] A. D. Rosenbloom, E. W. Kovar, D. R. Kovar, L. M. Loew, and T. D. Pollard, Biophysical Journal

**120**, 4399 (2021).

[11] D. Sept and J. A. McCammon, Biophysical journal **81**, 667 (2001).

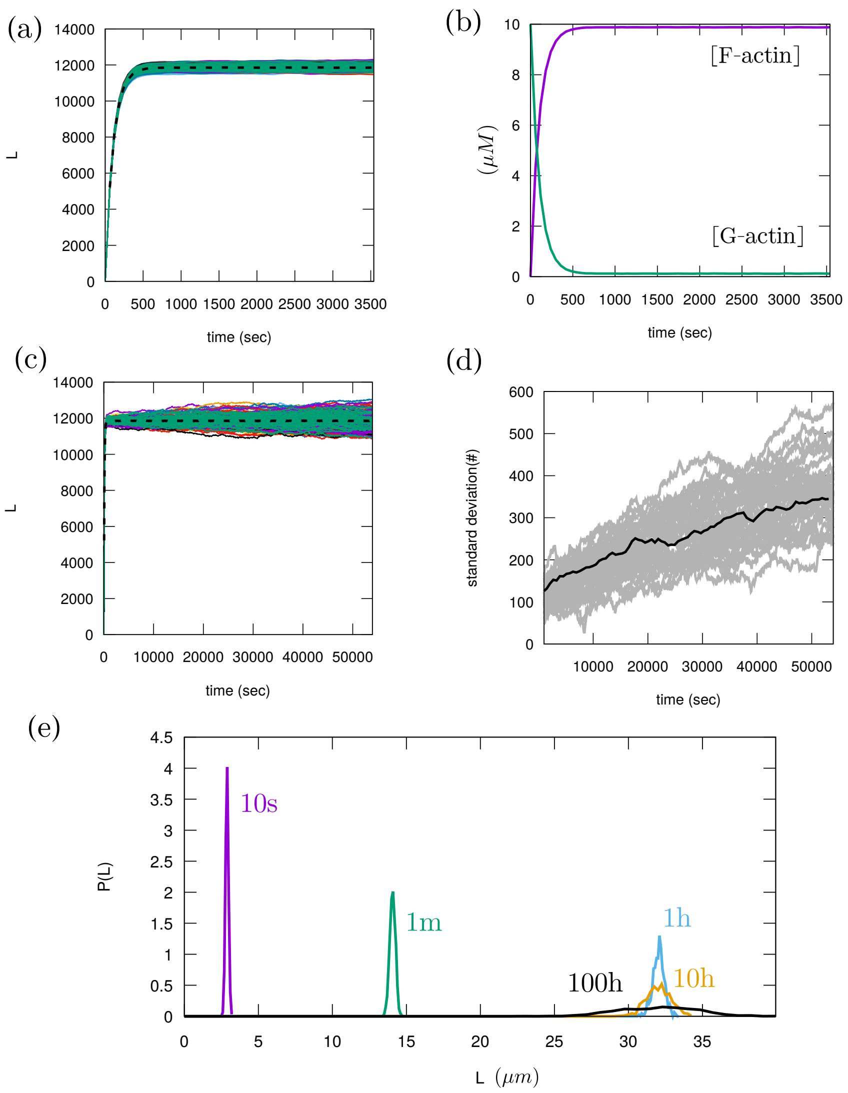


FIG. S1. **Early time convective and late time diffusive dynamics of filament length distribution in a limiting monomer pool**. (a) The mean filament length (black dashed line) quickly reaches equilibrium and remains unchanged. Simultaneously growing filaments (i.e., all nucleated at $t = 0$) have very small heterogeneity in length over a considerably long time (1 hr). (b) The filament population size (i.e., sum of all filament lengths) and monomers reach equilibrium at the end of the convective period. (c-d) The mean length (black dashed line) remains unchanged in the diffusive regime of growth while the standard deviation increases with time as the length distribution very slowly approaches towards an exponential steady state. Individual filament fluctuations arising from the lack of robust length control in a limiting pool contributes to the increase in the breadth of the length distribution. In panel a,c and d the filament length and standard deviation are given in monomer units. (e) Time evolution of filament length distribution for multiple filaments in a limiting pool without spontaneous nucleation (i.e., all nucleated at $t = 0$). The time evolution clearly captures the early time convective growth and the late time diffusive growth. We used 40 ensembles each with 10 filaments growing at total (monomeric and filamentous) actin concentration of 10$\mu$M. Only reaction 6 was used with growth rates as reported in the main text Table. 1.

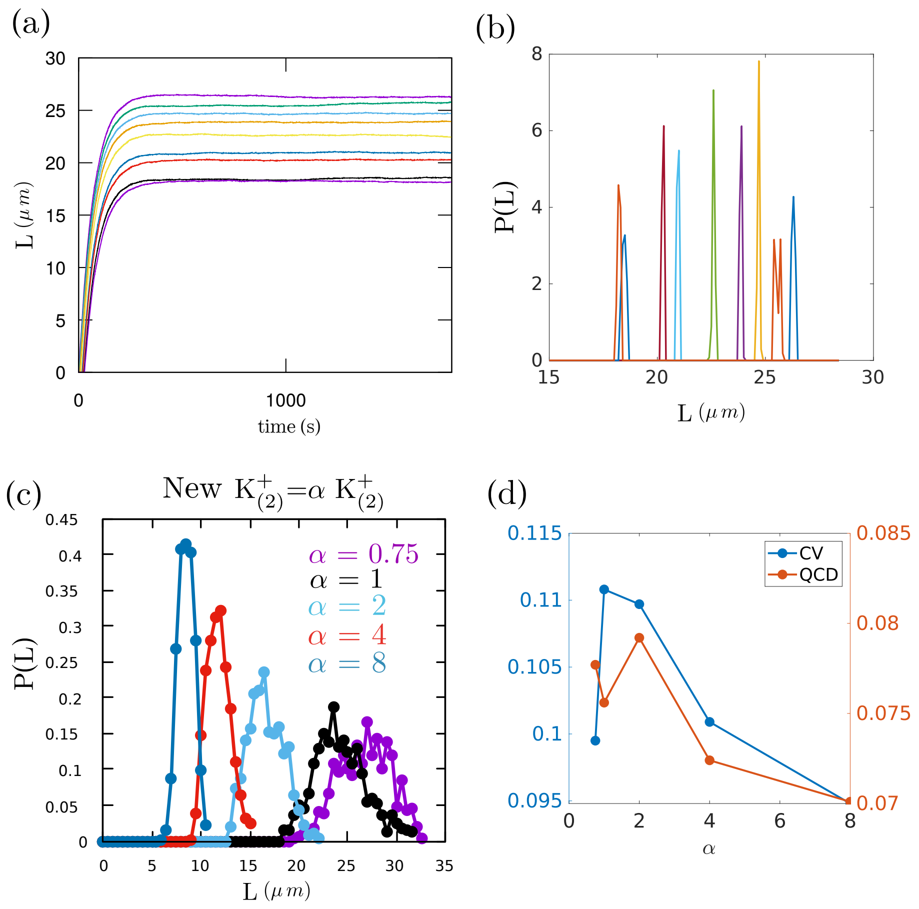


FIG. S2. **Actin filament length heterogeneity from spontaneous nucleation**. (a-b) Filament growth in time and the resulting length distribution (during last minute of simulated growth) show that the fluctuations in individual filament length are small. The heterogeneity in length arises due to sequential nucleation and not form individual filament length fluctuations. (c) The nucleation rate determines the amount of heterogeneity in length. Here we changed the dimer formation rate ($K^+_{(2)}$) to show that at faster nucleation rate (than spontaneous actin nucleation when $\alpha > 1$) the filament population has a narrower length distribution. (d) Quantification of the length heterogeneity by coefficient of variation (CV) and quartile coefficient of dispersion (QCD) captures the decrease in heterogeneity with increasing nucleation rate. We regard the non-monotonic trend as fluctuations arising due to limited number of ensemble average ($N = 20$). Parameter values used are same as in Fig. 2c with spontaneous nucleation.

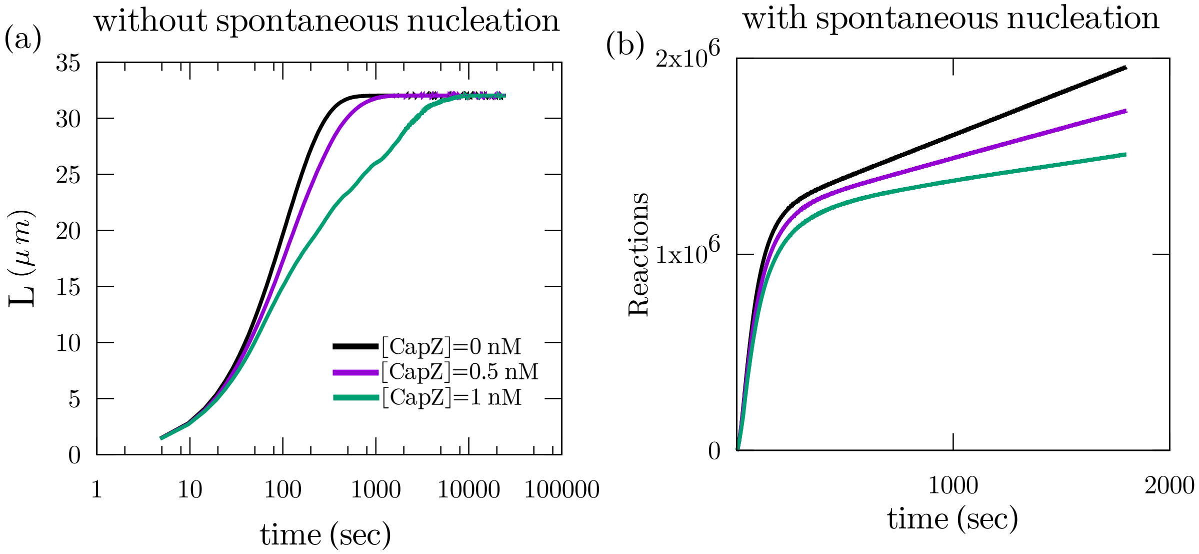


FIG. S3. **Effect of capping proteins on filament nucleation and diffusive growth**. (a) Presence of capping proteins cannot change the mean filament length at steady-state if spontaneous nucleation is not considered. The growth of the mean length becomes increasingly slower with increasing capping concentration but finally becomes same irrespective of the capping concentration. (b) Total number of polymerization reactions (sum of assembly and disassembly reactions) decreases with increasing capping concentration in the diffusive growth regime. This means that the length rearrangement in between the filaments is slower in presence of capping. Parameter values used for (a) are same as in Fig. S2 with 100 filaments growing in the system. Parameter values used for (b) are same as in Fig. 2c with spontaneous nucleation.

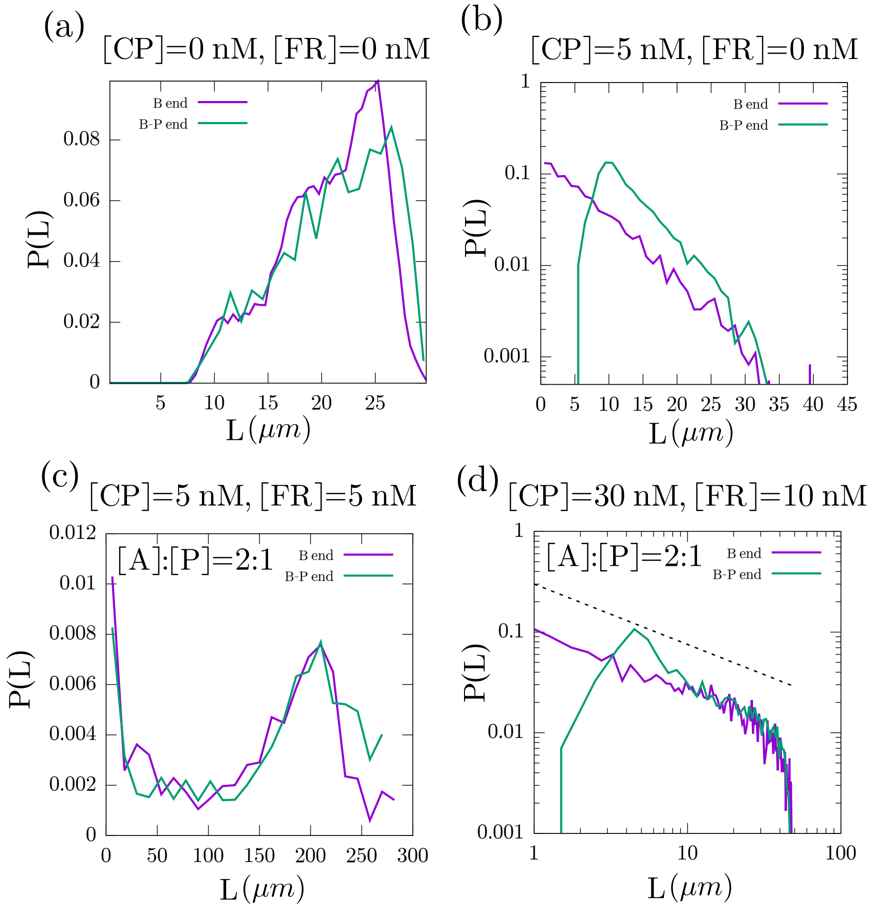


FIG. S4. **Effect of pointed end assembly on actin length distribution.** (a) In the absence of formin and capping proteins, the length distribution remains qualitatively similar witha very small increase in mean length due to excess polymerization in the pointed end. The legend "B end" denotes cases where the filament can only grow/shrink from the barbed end and the legend "B-P end" denotes cases where the filament can grow/shrink from both barbed and pointed end. (b) In the presence of capping proteins, the peak of the length distribution shifts to a larger value due to the pointed end assembly. The decay/tail of the length distribution remains exponential. (c) In the case of bimodal length distribution, there is no noticeable qualitative or quantitative change. (e) In the case of the fat tailed distribution, the peak of the length distribution shifts to a larger value due to the pointed end assembly but the tail of the length distribution shows a power-law decay. The parameter values used here are same as in Fig. 5 with $K_{FN} = 10^{-6} \mu M^{-2} s^{-1}$. The assembly and disassembly rates from pointed ends used are $1.3\,\mu\mathrm{M}^{-1} s^{-1}$ and $0.27\,s^{-1}$ respectively [9].

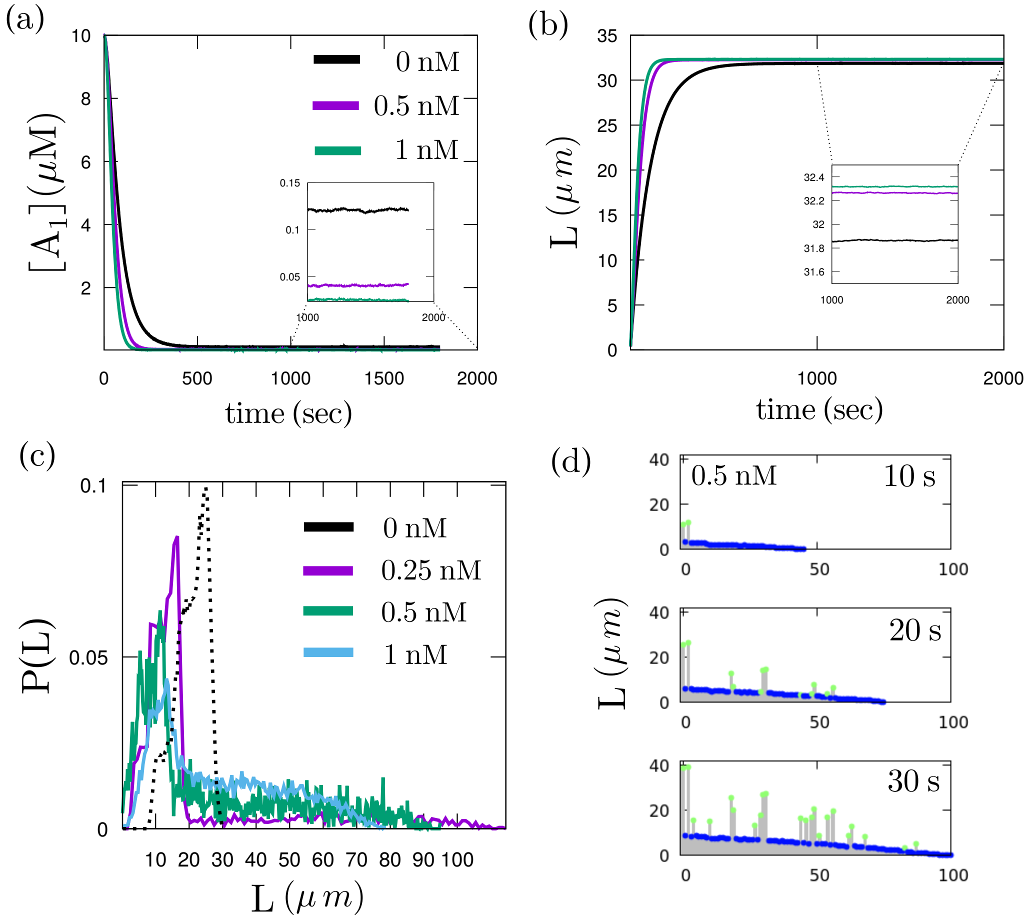


FIG. S5. **Effect of formin on filament length control and heterogeneity**. (a) In the presence of formin, the equilibrium concentration of the monomer pool depends on the abundance of formin and decreases with increasing formin concentration. The legends indicate the formin concentration. (inset) A zoomed-in view of filament length dynamics during the last 800 seconds of simulated growth. (b) The mean filament length increases with increasing formin concentration. (inset) A zoomed-in view of filament length dynamics during the last 1000 seconds of simulated growth. These results are obtained from simultaneous growth of all filaments without spontaneous nucleation. (c) Actin filament growth with spontaneous nucleation in the presence of formin proteins shows a state of increased length heterogeneity with length distributions exhibiting fat tails at low formin concentration. The legends indicate formin concentration value for the corresponding colour. (d) The origin of this length heterogeneity lies in the emergent competition between formin-bound filaments (marked by green dots) and filaments with free barbed end (marked by blue dots) for the monomers. This creates a subpopulation of large filaments that are formin-bound. This effect goes away at high formin concentrations as there are enough formin proteins to bind all the filaments. Parameter values used for (a-b) are same as in Fig. S3. Parameter values used for (c-d) are same as in Fig. 4 with the effective growth promotion by formin proteins.

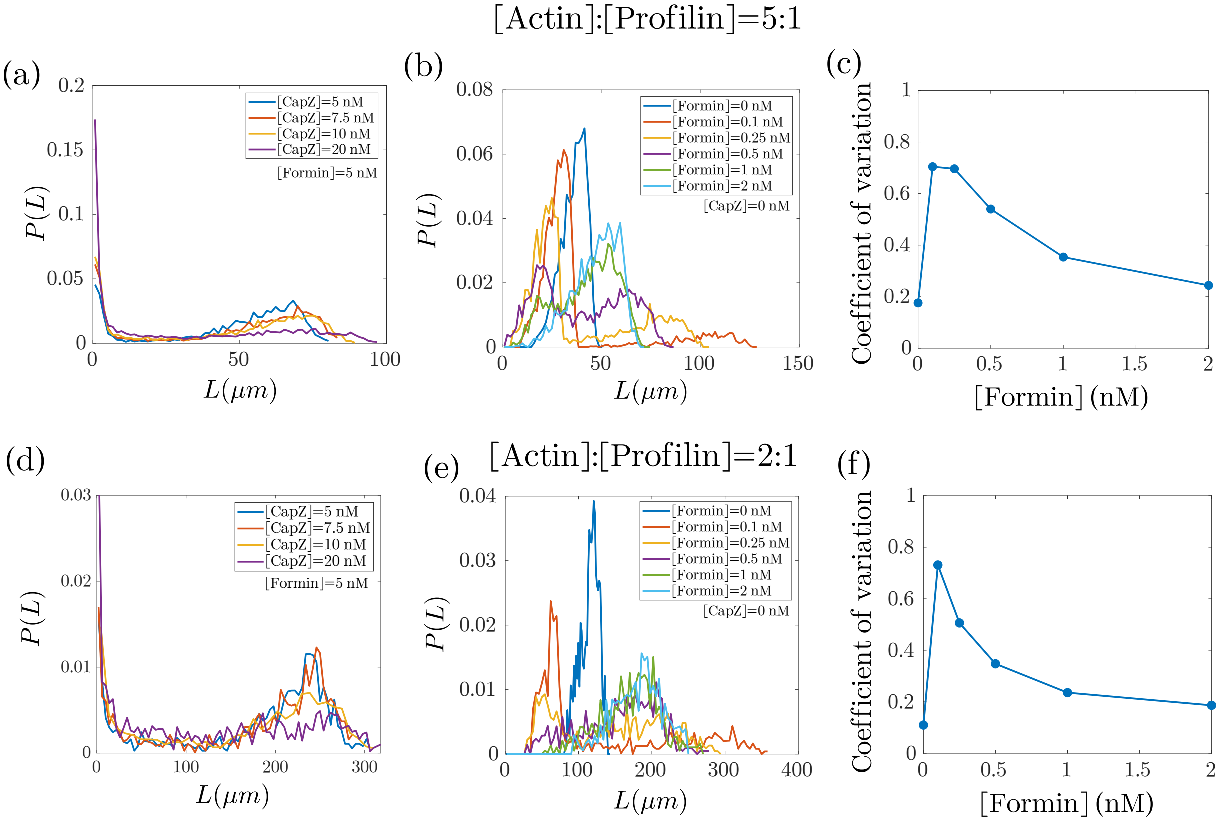


FIG. S6. **Effect of profilin on F-actin length regulation**. (a-c) [actin]:[profilin]=5:1. (a) In the presence of profilin, we find bimodal filament length distribution when capping and formin concentrations are comparable (blue, orange and yellow lines). The length distribution undergoes a transition to become fat-tailed at higher concentrations of capping proteins (magenta line). (b) Length distribution shows increased heterogeneity at low formin concentrations. (c) The coefficient of variation changes non-monotonically, consistent with the results shown in panel b. (d-f) The results remain qualitatively similar at a higher profilin concentration [actin]:[profilin]=2 : 1. (d) In the presence of profilin, we find bimodal length distribution when capping and formin concentrations are comparable (blue, orange and yellow lines). The length distribution becomes fat-tailed at higher capping concentrations (magenta line). (e) Length distribution shows increased heterogeneity at low formin concentrations. (f) The coefficient of variation changes non-monotonically, consistent with the results shown in panel e. Detailed description of the interaction between profilin, actin and formin is provided in Supplementary Sec **??**. Parameter values used here are same as Fig. 5 with $K_{FN} = 0 \mu M^{-2} s^{-1}$.

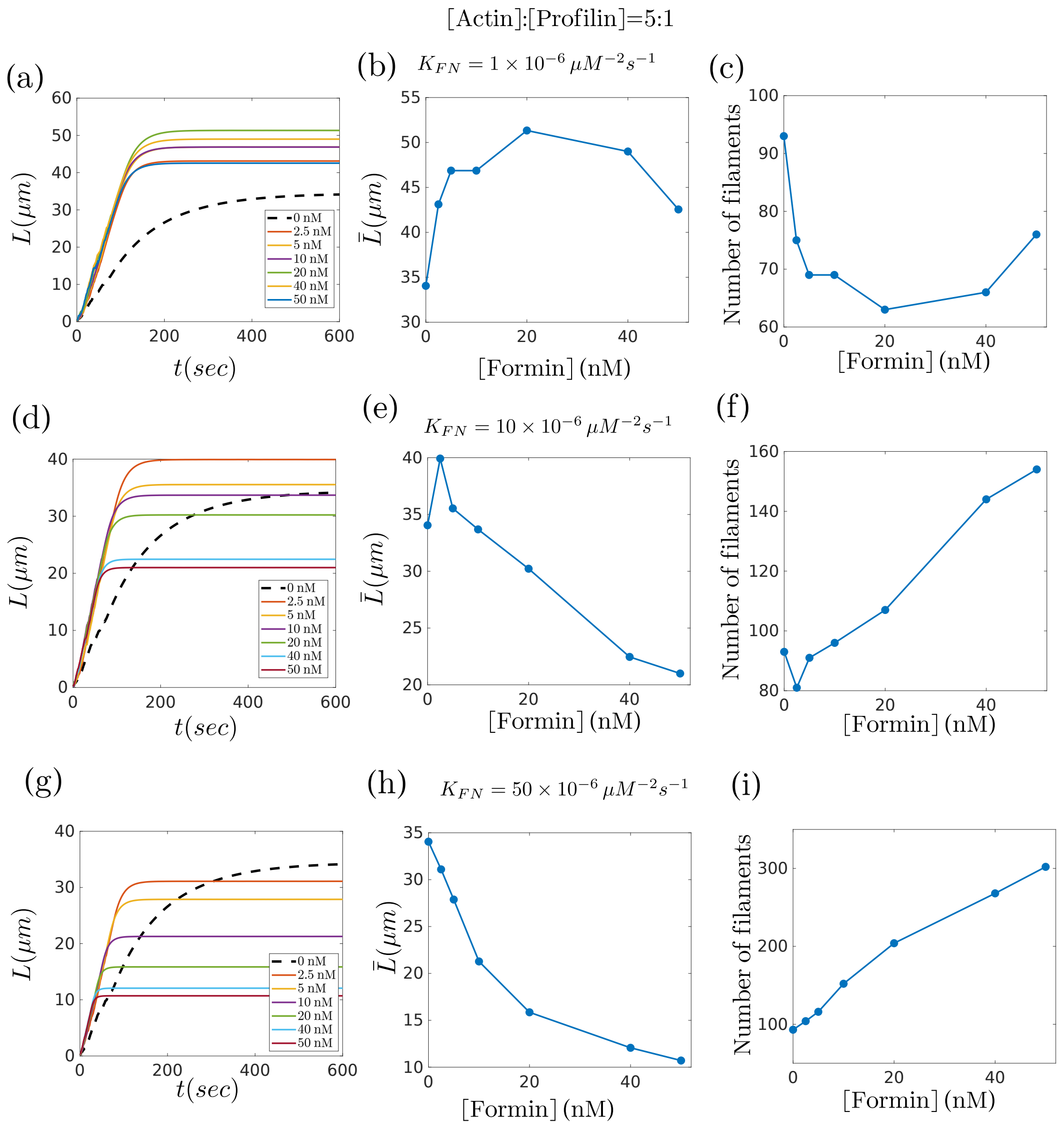


FIG. S7. **Effects of formin mediated filament nucleation on F-actin length regulation for [actin]:[profilin]=5:1**. (a-c) For $K_{FN} = 10^{-6} \mu M^{-2} s^{-1}$. (a) Mean filament length vs time for different formin concentrations. The legend indicates the formin concentrations. (b) The mean filament length changes non-monotonically with formin concentration. (c) Filament number decreases with formin concentration at smaller concentrations of formins but increases at larger formin concentration. (d-f) For $K_{FN} = 10^{-5} \mu M^{-2} s^{-1}$. (d) Mean filament length vs time for different formin concentrations. (e) The mean length changes non-monotonically with formin concentration. (f) Filament number decreases with formin concentration at very small concentrations of formins but increases at larger formin concentration. (g-i) For $K_{FN} = 5 \times 10^{-5} \mu M^{-2} s^{-1}$. (g) Mean filament length vs time for different formin concentrations. The legend indicates the formin concentrations. (h) The mean length decreases with increasing formin concentration. (c) The filament number increases monotonically with formin concentration. For details of the formin-meadiated nucleation process see Supplementary Sec **??**. Reactions. $1 - 14$, described in Fig. 1, were used to obtain these results with the rates mentioned in main text Table. 1. Profilin concentration of $2\mu$M was used with all other parameter values are same as in Fig. 5.

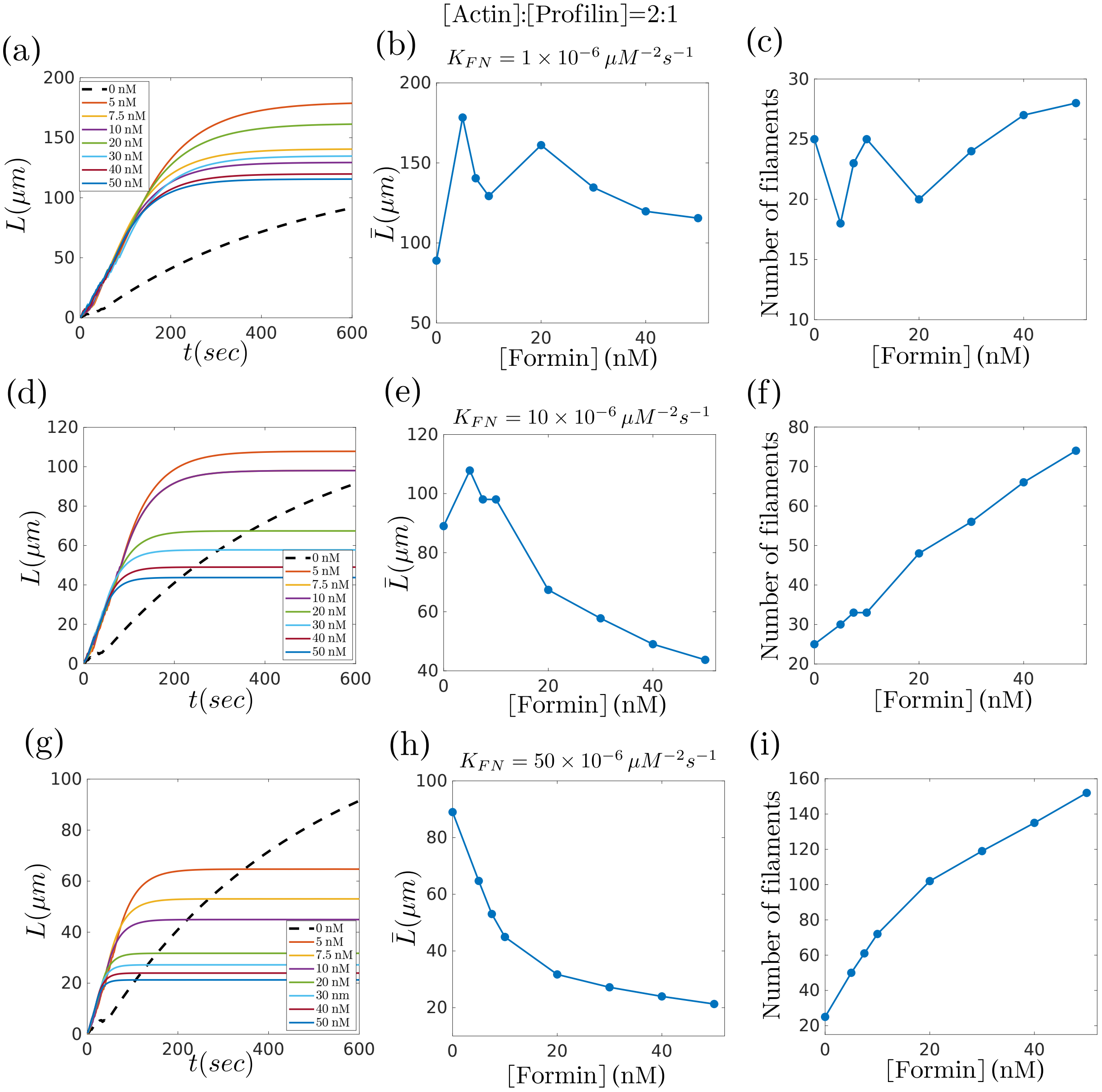


FIG. S8. **Effects of formin mediated filament nucleation on F-actin length regulation for [actin]:[profilin]=2:1**. (a-c) For $K_{FN} = 10^{-6}\mu M^{-2}s^{-1}$. (a) Mean filament length vs time for different formin concentrations. The legend indicates the formin concentrations. (b) The mean filament length changes non-monotonically with formin concentration. (c) Filament number decreases with formin concentration at smaller concentrations of formins but increases at larger formin concentration. (d-f) For $K_{FN} = 10^{-5}\mu M^{-2}s^{-1}$. (d) Mean filament length vs time for different formin concentrations. (e) The mean length changes non-monotonically with formin concentration. (f) Filament number increases with formin concentration. (g-i) For $K_{FN} = 5 \times 10^{-5}\mu M^{-2}s^{-1}$. (g) Mean filament length vs time for different formin concentrations. The legend indicates the formin concentrations. (h) The mean length decreases with increasing formin concentration. (c) The filament number increases monotonically with formin concentration. For details of the formin-meadiated nucleation process see Supplementary Sec **??**. Reactions. $1 - 14$, described in Fig. 1, were used to obtain these results with the rates mentioned in main text Table. 1. Profilin concentration of $5\mu$M was used with all other parameter values are same as in Fig. 5.

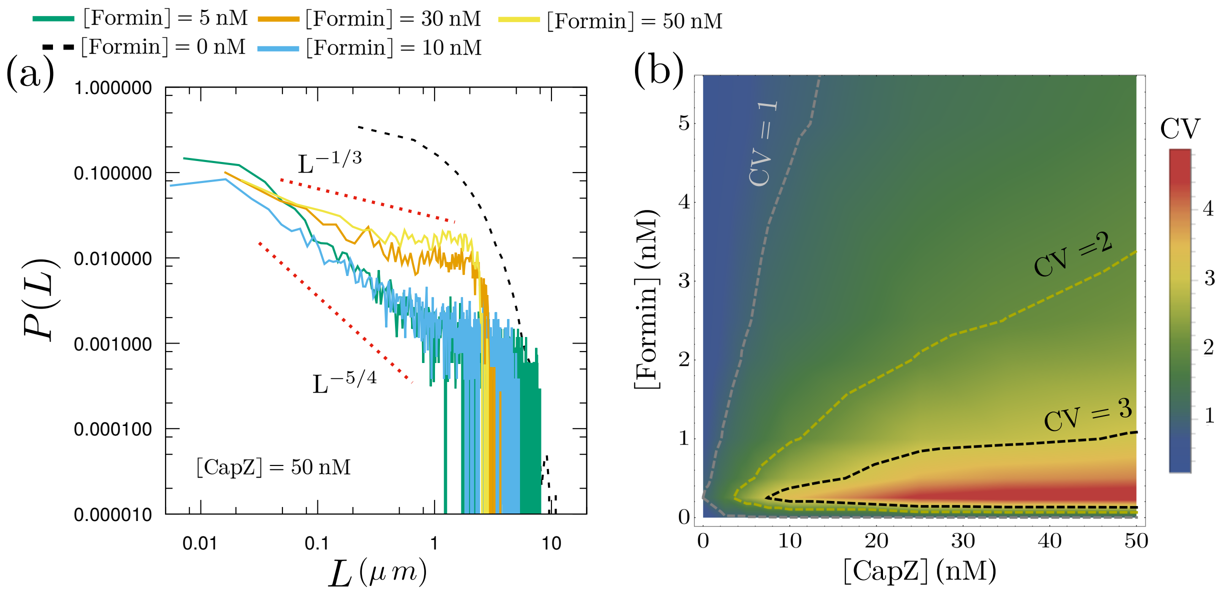


FIG. S9. **The fat-tailed phase exhibits power law decay in filament length distribution**. (a) Filament length distribution in the the fat-tailed phase shows power law decay followed by a terminal exponential decay. The exponent of the power law decreases with increasing formin concentration. This feature arises in a large parameter region between the exponential phase at very low formin concentration and the bimodal phase at higher formin concentration. (b) A zoomed in phase diagram of the coefficient of variation (CV) as a function of capping and formin concentration. The fat-tailed phase exhibits high amount of length heterogeneity. The contours trace the iso-CV lines. Parameter values used here are same as in Fig. 6

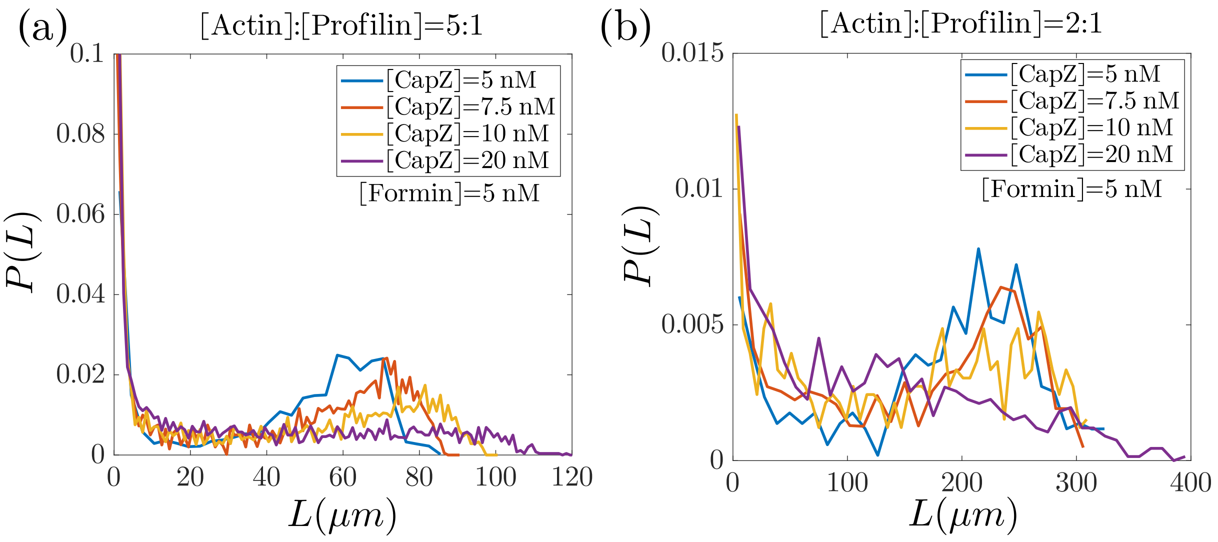


FIG. S10. **Bimodal and fat-tailed length distribution in presence of formin-mediated filament nucleation.** (a-b) In the presence of formin-mediated nucleation, we find bimodal length distribution when capping and formin concentrations are comparable (blue and orange lines). The length distribution becomes fat-tailed at higher capping concentrations (magenta line). The dimer formation rate was taken to be low ($K_{FN} = 10^{-6}\mu M^{-2}s^{-1}$) corresponding to formins that are inefficient nucleators without nucleation promoting factors (e.g., mDia1). For additional details of the formin and profilin reactions see Supplementary Sec **??**. All reactions described in Fig. 1 were used to obtain these results with the rates mentioned in main text Table. 1. Profilin concentration of 2$\mu$M(a) and 5$\mu$M(b) were used with all other parameter values are same as in Fig. 5.

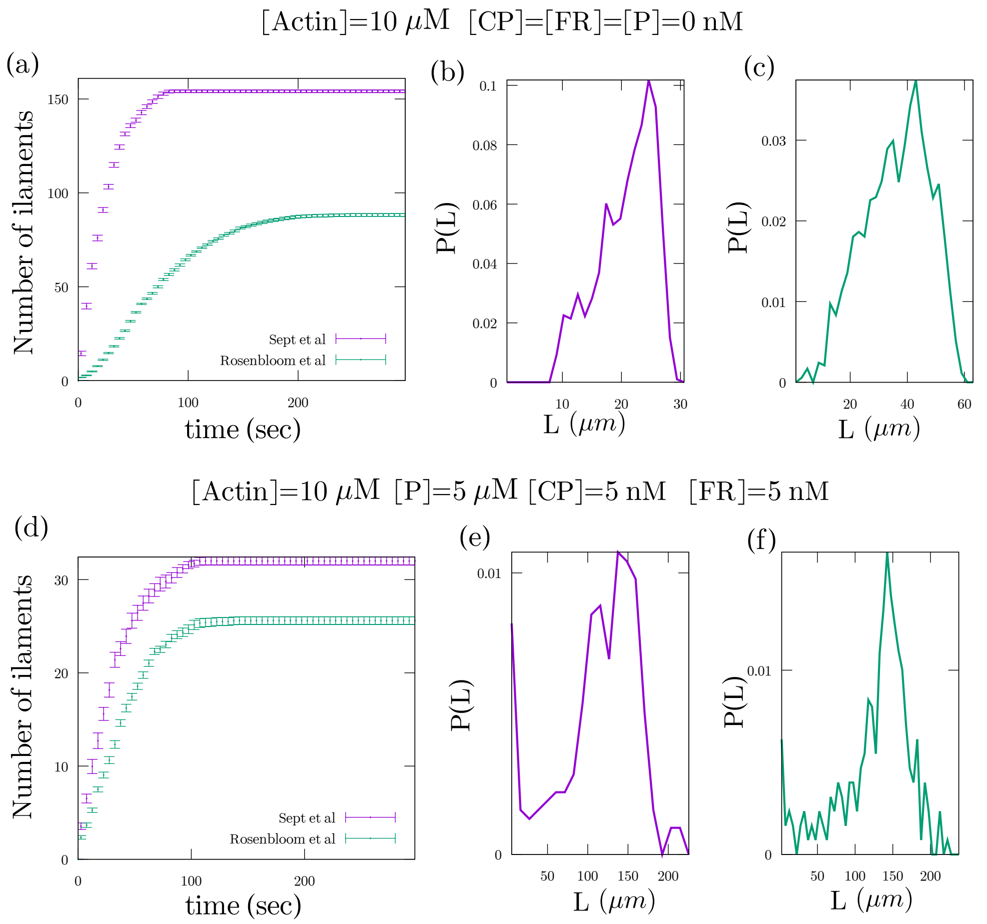


FIG. S11. **Effect of nucleation rates on filament growth and nucleation rates.** (a-c) Case-I with only actin: (a) Less number of filaments are slowly nucleated when using nucleation rates from Rosenbloom *et al* [10] compared to the rates reported by Sept and McCammon [11]. (b-c) The length distributions are qualitatively similar with a higher mean length with the new nucleation rates (right). (d-f) Case-II with actin, capping, formin and profilin: (d) The filament number is smaller with the new nucleation rate. (e-f) The length distributions obtained using the two different nucleation rates are qualitatively similar with small quantitative difference. There is a prevalence of larger filaments with the new slower nucleation rates reported in Rosenbloom *et al* (right). The values of the new dimer and trimer formation and dissociation rates are: $K^{+}_{(2)} = 3.5 \times 10^{-6}\,\mu\text{M}^{-1}s^{-1}$, $K^{-}_{(2)} = 0.041\,s^{-1}$, $K^{+}_{(3)} = 13 \times 10^{-5}\,\mu\text{M}^{-1}s^{-1}$, $K^{-}_{(3)} = 22\,s^{-1}$ as obtained in Rosenbloom *et al* [10]. Here $K_{\text{FN}} = 10^{-5}\,\mu M^{-2}s^{-1}$ and all other rates are same as mentioned in the parameter tables in main text.

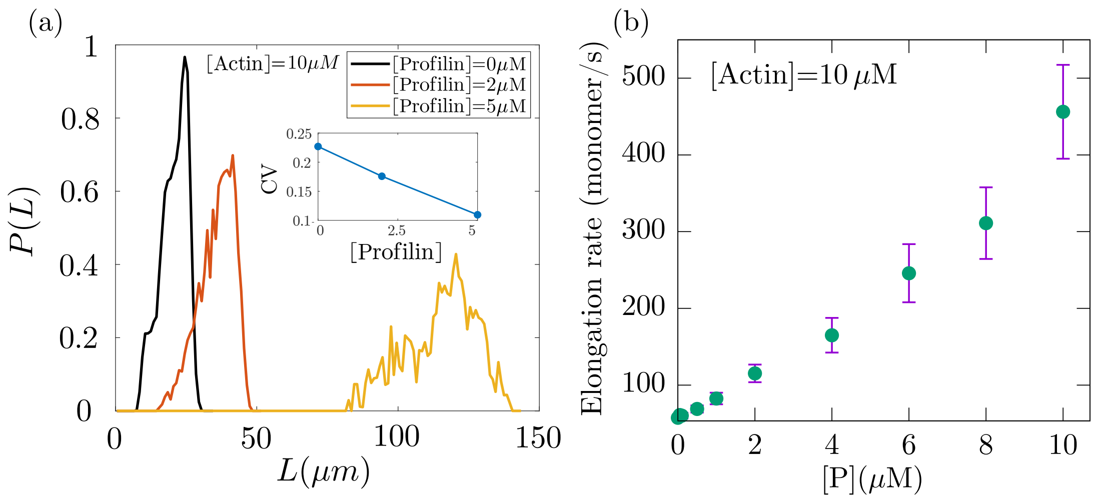


FIG. S12. **Profilin decreases filament length heterogeneity and increases elongation rate.**(a) In the absence of formin and capping proteins, profilin-induced reduction in spontaneous nucleation leads to lower filament density. This results in increased mean length and smaller heterogeneity. (inset) Quantification of the coefficient of variation (CV) shows a reduction in value with increasing profilin density. (b) Barbed end elongation rate, calculated from the first 50 seconds of filament growth, increases with increasing profilin concentration. The formin concentration used in panel-b is 5 nM. For additional details of profilin and formin reactions see Supplementary Sec **??**. Parameter values used here are the same as in Fig. 5.
